# Identification of Therapeutic Leads from *Ficus hispida* Fruit Phytochemicals against Prostate Cancer Using Pharmacoinformatic and Molecular Dynamics Simulation Approach

**DOI:** 10.1101/2023.06.04.543596

**Authors:** MD. Hasanur Rahman, Md. Ataur Rahman, Md. Sarwar Zahan, Partha Biswas, Silme Islam, Riyan Al Islam Reshad, Abdullah Al Mamun Sohag, Bablu Gupta, Redwanul Islam, Md. Abdul Hannan, Woojin Kim, Seungjoon Moon, Md Jamal Uddin, Bonglee Kim

## Abstract

Prostate cancer is one of the leading causes of death and the most common cancer type in men. In this study, potential leads from the phytochemicals of *Ficus hispida* fruit were screened using in silico tools against androgen receptor (AR), a known target for prostate cancer. PASS online and ADMET tools were used to screen specific phytochemicals that are relevant to prostate cancer treatment and have drug-like properties. Of 13, a total of 10 phytochemicals passed PASS online and ADMET screening. Next, a total of three phytochemicals, including nodakenetin (CID: 26305), isowigtheone hydrate (CID: 66728267), and 7-hydroxycoumarin (CID: 5281426) were selected based on their docking scores (-9.946 to -7.653 kcal/mol) and relevance to selective bioactivity. The MD simulation further confirmed the binding stability of these three phytochemicals with their target AR protein and determined that the main amino acid residues mainly responsible for this stability including RMSD, RMSF, and post simulation binding free energies. These findings suggest that these three phytochemicals of *Ficus hispida* fruit can further be developed as prospective therapeutics against prostate cancer.

## 1. Introduction

Prostate cancer is the most common cancer in men and the second leading cause of death, accounting for 1.41 million global deaths in prostate cancer ^1^, with over 160,000 new cases diagnosed annually in the United States ^2^. Despite its high prevalence, prostate cancer represents a clinically diverse spectrum of disease, with some patients exhibiting aggressive outcomes while others show a low tendency to progress. The high mortality rate is partially attributed to the lack of early detection and effective treatment options for curbing metastasis. Standard treatments for prostate cancer include radiotherapy and surgery, while patients unsuited to these options are treated with androgen ablation therapy. However, this treatment is often followed by recurrent androgen-independent prostate cancer with frequent metastasis, significantly impacting the patient’s quality of life ^3–5^. Androgen receptor (AR) belongs to a group of nuclear steroid hormone receptors and is crucial for the development of the prostate glands. Physiologically, it regulates gene transcription, cell division and death and is crucial for the survival and growth of prostate cells, making it a prime target for prostate cancer treatment ^6–8^. The growth of prostate tumors is largely dependent on the androgen receptor pathway, therefore blocking this pathway is a useful approach to managing prostate cancer ^9^.

Studies have shown that consuming vegetables and fruits high in phytochemicals can decrease cancer risk ^10^. *Ficus hispida* fruits are rich in polyphenolics and have demonstrated various health benefits, including anticancer potential ^11^. Evidence shows that the methanolic extract of *Ficus hispida* fruits has cytotoxic effects on certain human cancer cell lines, and the leaf and bark extracts have growth inhibitory effects ^12–14^. However, the potential lead phytochemicals of *Ficus hispida* fruits and the specific mechanisms they employed to inhibit the growth of cancer cells are yet to be explored.

The purpose of this study was to identify therapeutic leads from *Ficus hispida* fruit phytochemicals against AR for the prevention of prostate cancer. And the ultimate goal is to develop new drugs based on these phytochemicals that could be used for the prevention and treatment of prostate cancer. To achieve these goals, various computational approaches, including ADMET screening, molecular docking and dynamics simulation were exploited as a preliminary tool for drug discovery.

In this study, potential anticancer phytochemicals were initially selected from the *Ficus hispida* fruit chemicals based on their pharmacokinetic properties, including ADMET ^15^. The best binding ligands were then selected to construct on their anticancer properties through PASS online results and molecular docking studies against AR protein (PDB ID: 5T8E). To validate the results from the docking study, molecular dynamics simulations and MM-GBSA were conducted using the Desmond ^16^ software. This computational pipeline exploited various computational methods for the design of pharmaceuticals from *Ficus hispida* fruit phytochemicals aiming at the development of the most promising therapeutic candidates for use in the effective treatment of prostate cancer.

## 2. Materials and Methods

### 2.1 Ligand Library Generation

Dr. Duke’s phytochemical and ethnobotanical database (https://phytochem.nal.usda.gov/phytochem/search), as well as reviews and research articles were used to generate a library of the phytochemicals for this study ^12, 17^. A total of 89 phytochemicals of *Ficus hispida* were found in these sources (Supplementary Table-1). Since this study focuses on the fruit part, only 13 anti-prostate cancer phytochemicals (found in fruit) were chosen (Table-1). The selected phytochemical structures were available and retrieved from the PubChem database at the National Institutes of Health website (https://pubchem.ncbi.nlm.nih.gov/) ^18^.

### 2.2 Ligand Activity Prediction

A popular server named PASS (Prediction of Activity Spectra for Substances) (http://www.way2drug.com/passonline/), which predicts outcomes based on a substance’s structural constitution, was used to compare the biological activity of the selected 13 fruits phytochemicals ^19^. The Structure Activity Relationship Base is used to predict the likelihood that a specific substance will belong to the active and inactive subsets of that substance (SAR Base). The input was provided in SMILES (Simplified Molecular Input Line Entry System) format for the phytochemicals structure. We calculated the Pa (probable activity) and Pi (probable inactivity) values for each ligand. Therefore, only the activities with cancer and androgen antagonists were taken into consideration ^20^.

### 2.3 ADMET Studies

In clinical trials, QikProp ^21^ finds lead compounds that perform better in terms of ADMET (absorption, distribution, metabolism, excretion, and toxicity) ^22^. QikProp is a powerful ADMET prediction tool for pharmaceutical analysis. We further analyzed the past 10 bioactive ligand molecules from PASS analysis with pharmacological properties linked to prostate cancer (Table-2).

### 2.4 Protein Retrieval and Preparation

This study aimed to develop prostate cancer androgen receptor modulators with improved pharmacokinetic profiles and metabolic stability. After searching for the protein structures of androgen receptors associated with prostate gland, in RCSB PDB protein data bank (www.rcsb.org) we found the PDB ID: 5T8E as an ideal candidate for this study which has a suitable PK profile and improved metabolic stability while maintaining potent androgen agonistic activity. It provides a high-resolution model of the androgen receptor ligand-binding domain, which is the primary target of androgen agonists ^23^.

However, the X-ray crystallographic structure of Androgen receptor protein (PDB ID: 5T8E) was retrieved. To perform better interactions with selected ligands to a targeted protein, the protein crystal structure must prepare well. Therefore, protein crystal structures need to be built before docking so that hydrogen atoms can be added, hydrogen bonds can be optimized, atomic conflicts can be reduced, and various other tasks can be executed that are not part of the x-ray crystal structure refinement process ^24^. The protein preparation wizard in Schrödinger suite 21.2 was used to prepare the retrieved 5T8E protein structure. In the protein preparation wizard, assigned bond orders, CCD database, the addition of hydrogens, the formation of zero-order bonds to metals, and the formation of disulfide bond properties were used. Moreover, to fix the protein missing side chain and loops in the receptor, the missing side chains, and loops properties were used by using Prime in Schrödinger suite 21.2 ^25^. Also, Epik in Schrödinger suite 21.2 ^26^ were used to calculate and fix cap termini, waters of beyond 5Å, and the generation of heat states pH level 7.0 +/- 2.0. The refine tab and the OPLS3e force field were used to assign the H-bond in PROPKA at pH level 7.0 and restrict the degradation to RMSD 0.30 Å with heavy atom coverage ^27^.

### 2.5 Ligand Preparation

The standardized drugs and phytochemicals, three-dimensional (3D) structures were retrieved in SDF format with individual PubChem CID from the open-source PubChem database (www.pubchem.ncbi.nlm.nih.gov). All the ligand structures were prepared using LigPrep application in Schrödinger suite 21.2 ^28^. The OPLS3e force field and Epik ionizer were used to execute minimization at a standard pH range of 7.0 to (+/-) 2.0, with a maximum number of conformers per structure was 32 and an RMSD of 1.0 Å.

### 2.6 Receptor Grid Generation

The grid restricts the active site to a more condensed region of the receptor protein, and the grid makes it easier for the ligand to bind in its desired location ^29^. Initially, the Receptor Grid Generation tool included in Schrodinger suite 21.2 was utilized to construct a grid with a Van der Waals residue using the parameters of scaling factor = 1.0 and partial charge cutoff = 0.25. The co-crystalized inhibitor 77U of 5T8E protein (orange color figure 1) was used as a reference binding location for all the selected ligands and a cube-shaped enclosure was formed around the 77U active site (reference ligand active site). Following that, the volume of the grid box was modified such that it measured 14 x 14 x 14 Å degrees for the docking process.

**Figure 1:**
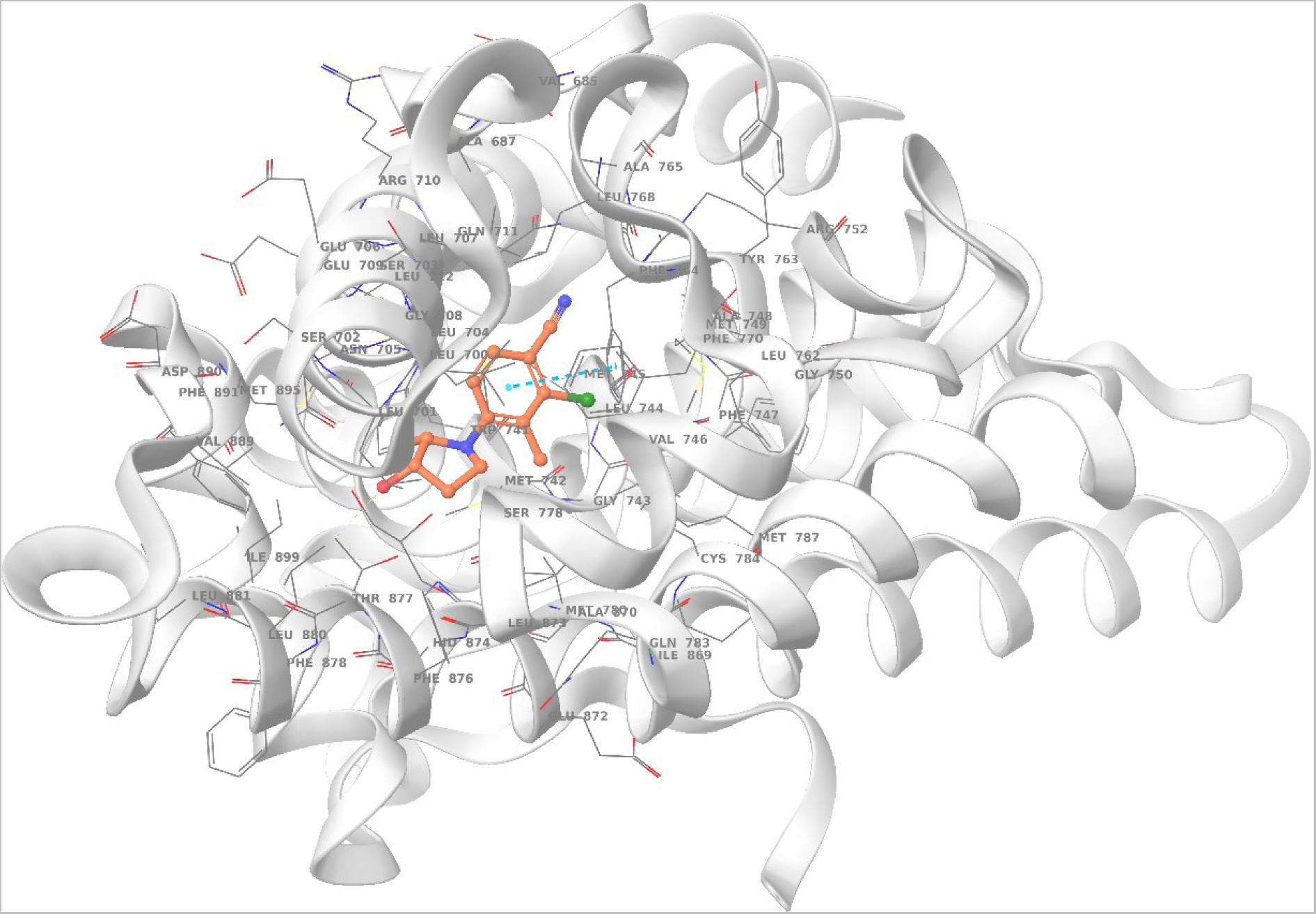
5T8E androgen receptor protein with associated 77U co-crystalized receptor (orange color) structure

### 2.7 Molecular Docking

The selected protein-ligand complexes structure was taken from a docked post-viewing file for post docking visualization purposes. Further, the study of non-bond interactions, hydrophobicity, and binding affinities of the complexes was evaluated by Maestro’s XP (extra precision) molecular docking application ^27^. The post-docked structures of the receptor-ligand complex were visualized using the Discovery Studio Visualizer 21 (BIOVIA, 2021). To clarify and visualize the non-boned and non-covalent bonded interactions, high binding affinities showed complexes were visualized in Discovery Studio Visualizer 21.

### 2.8 Molecular Dynamics (MD) Simulation

100 ns MD simulations were used to assess the consistency of the potential ligand phytochemicals binding to the targeted receptor active binding site. The thermodynamic stability of receptor-ligand complexes was examined using the Desmond application in Schrödinger suite 21.2 ^16^. Also, A predetermined TIP3P water technique was developed for this framework to maintain a specific volume with an orthorhombic periodic bounding box shape at a 10 Å distance. For electrically neutralizing the framework, suitable ions O+ and 0.15 M NaCl as salt have been selected and dispersed randomly inside the solvent system. After developing the solvency protein system with a ligand complex, the proposed system was optimized and simplified by applying the Desmond default protocols implemented by force field OPLS3e within the Desmond module ^30^. Moreover, A 50 ps capture period and an energy of 1.2 ps were utilized with the Nose-Hoover temperature combining technique and isotropic technique NPT. Then the final assemblies were maintained at 300 K and one atmospheric pressure (1.01325 bar). Finally, the results data of the 100 ns molecular dynamic simulation were visualized by using Simulation Trajectory Analysis (SID) application of Schrödinger’s suite 21.2 ^16^. SID was used to evaluate the stability of 100 ns runtime trajectory complex structures by root-mean-square deviation (RMSD), root-mean-square fluctuation (RMSF), radius of gyration (Rg), solvent-accessible surface area (SASA), protein-ligand contacts (P-L contact), ligand-protein contacts (L-P contact), and hydrogen bond interactions.

### 2.9 Post Simulation Binding Energy Calculation (MM/GBSA)

To estimate the free energies of the complexes during the 100 ns simulation we further calculated the MMGBSA by thermal_mmgbsa.py python package. The Desmond MD trajectory was split into 20 individual frame snapshots and runs each one through MMGBSA and then separated the ligand from the receptor.

## 3. Research Findings

### 3.1 PASS Online Prediction for QSAR Analysis

Using the PASS online tool, all 13 phytochemicals were assessed for their potential to prevent prostate cancer (Table 1). The phytochemicals were only taken into consideration based on the combined properties such as anti-metastatic, prostate cancer treatment, and androgen antagonist. Activities with greater Pa have pharmacological potency and experimental production potential^31^. But the activities selected in this study didn’t show a higher Pa value. Therefore, only the androgen antagonist was considered in eliminating the phytochemicals. Also, PASS can help to reduce the side effects of a molecule even though it cannot predict the binding affinity for new therapeutic targets. The filtered 10 phytochemicals mentioned above were taken for ADME analysis before being used in docking analysis.

**Table 1:**
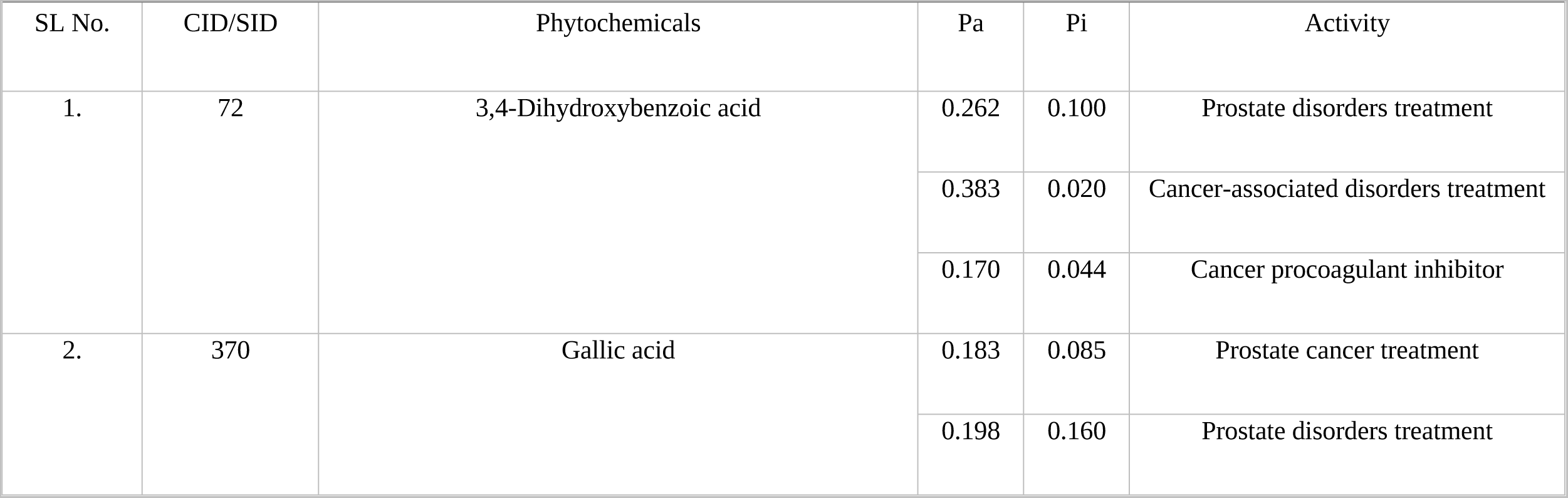
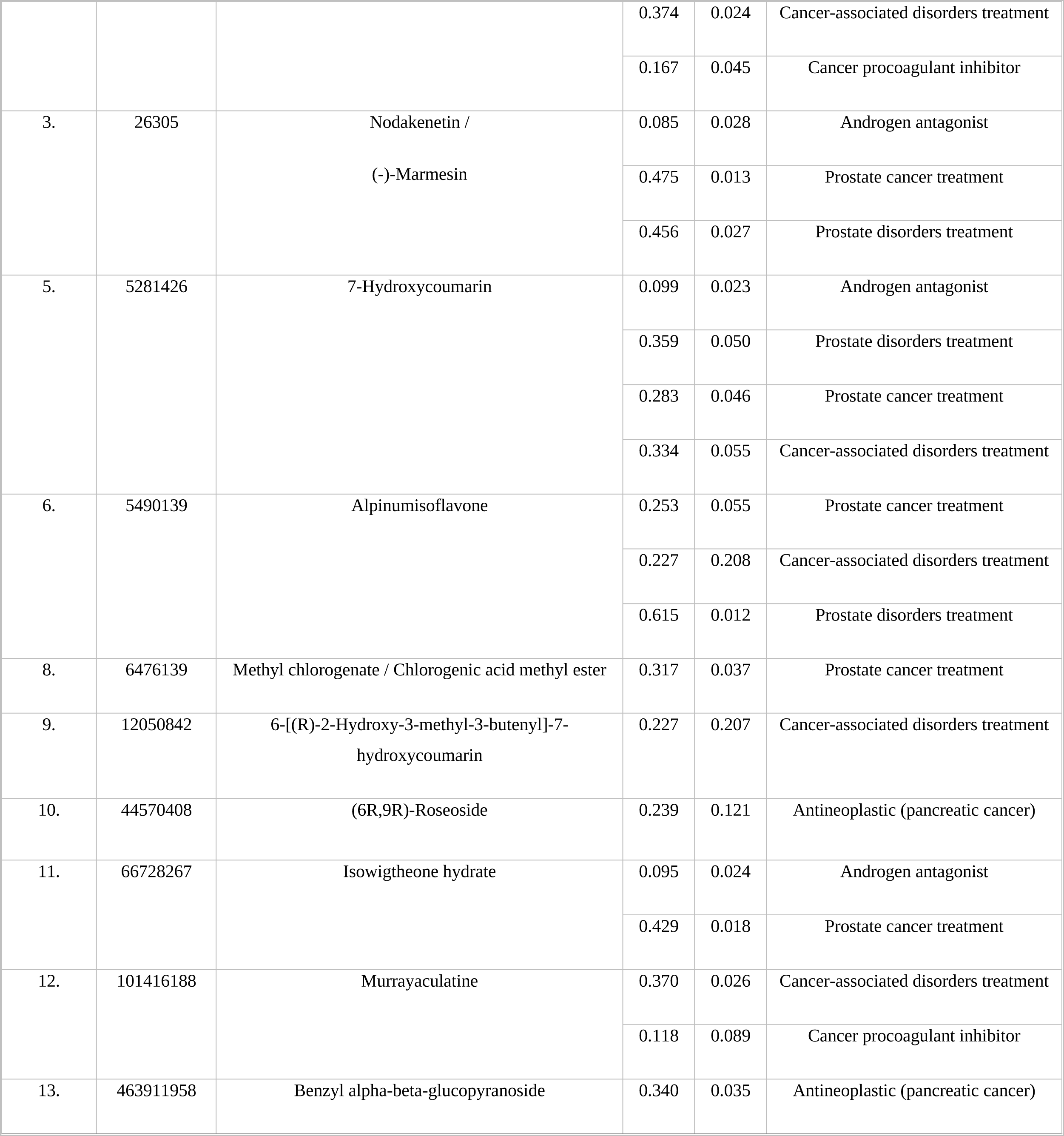
The result of QSAR models for bioactivity prediction in ligand validation.

### 3.2 ADMET Analysis

The ADMET properties of a drug candidate are crucial for the regulatory approval process since they assist researchers and drug developers to understand the safety and efficacy of a drug candidate ^32^. In this study, the phytochemicals that had been screened using the PASS program were assessed by QikProp for drug-like properties (ADME) investigations. For estimating the ADMET properties, variables such as molecular weight, molecular volume, QPlogPo/w, QPlogPw, QPlogS, QPlogKp, QPlogKhsa, QPCaco, and SASA were considered, whereas all the 10 phytochemicals undergone ADME screening and passed Lipinski’s filter. The chosen phytochemicals for the study fit within acceptable ranges, according to the analysis of ADMET qualities, showing their potential as drug-like phytochemicals (Table 2). The ADMET passed phytochemicals molecular weights range is, from 154.122 to 456.707. Also, the ADMET passed phytochemicals molar volume range is from 502.685 to 1387.897. 3,4-Dihydroxybenzoic acid (CID: 72), Gallic acid (CID: 370), Nodakenetin/(-)-Marmesin (CID: 26305), 7-Dihydroxycoumarin (CID: 5281426), Alpinumisoflavone (CID: 5490139), Methyl chlorogenate/Chlorogenic acid methyl ester (CID: 6476139), 6-[(R)-2-Hydroxy-3-methyl-3-butenyl]-7-hydroxycoumarin (CID: 12050842), (6R,9R) -Roseoside (CID: 44570408), Isowigtheone hydrate (CID: 66728267), and Murrayaculatine (CID: 101416188) demonstrated a complete adherence to Lipinski’s rule and was projected by QikProp analysis to have a high proportion of human absorption (Table 2). It is anticipated that phytochemicals that have passed Lipinski’s rules and the ADMET qualities will make for suitable oral medications ^33^.

**Table 2:**
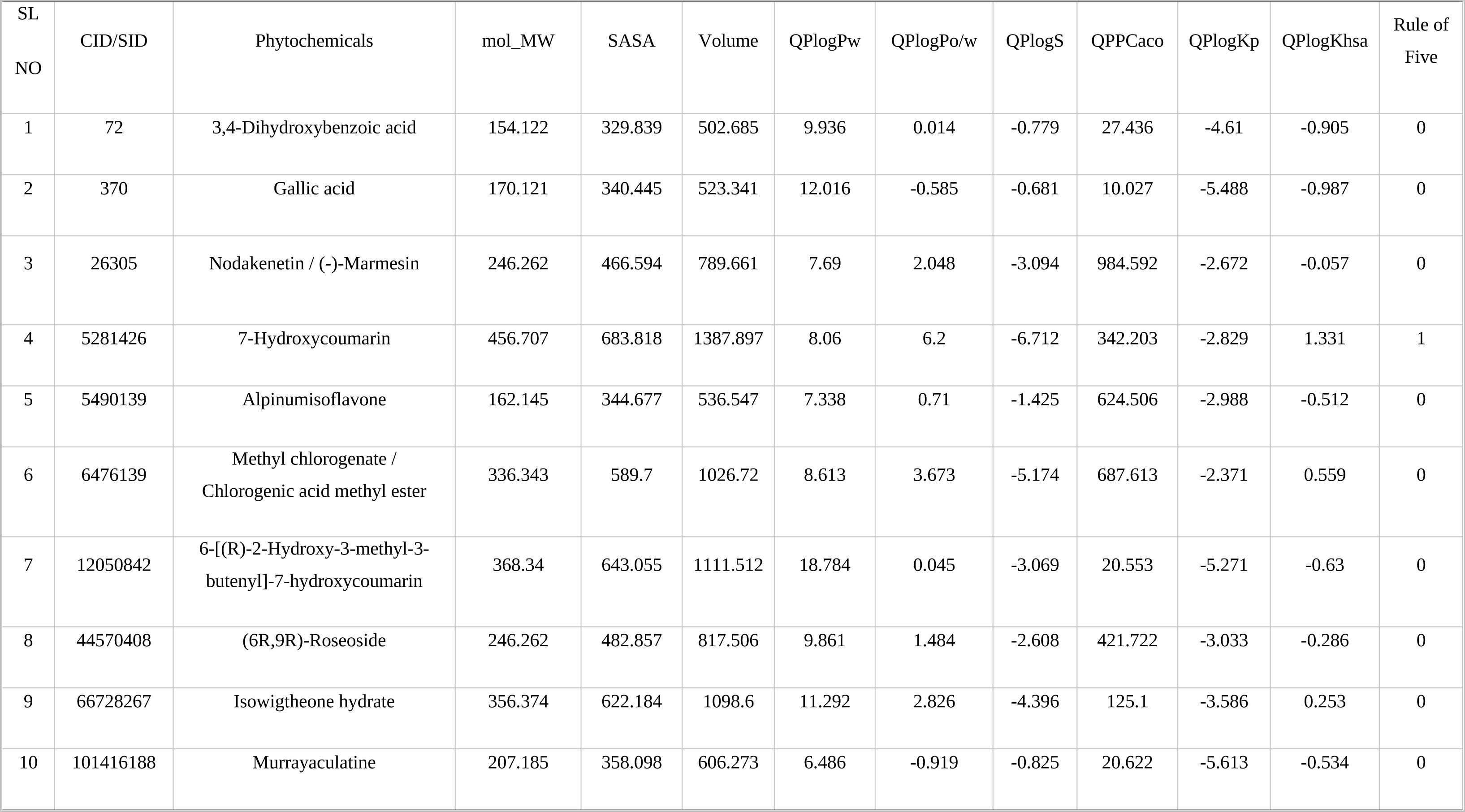
List of pharmacokinetics properties (ADMET) of all the selected phytocompounds.

**Table 3:**
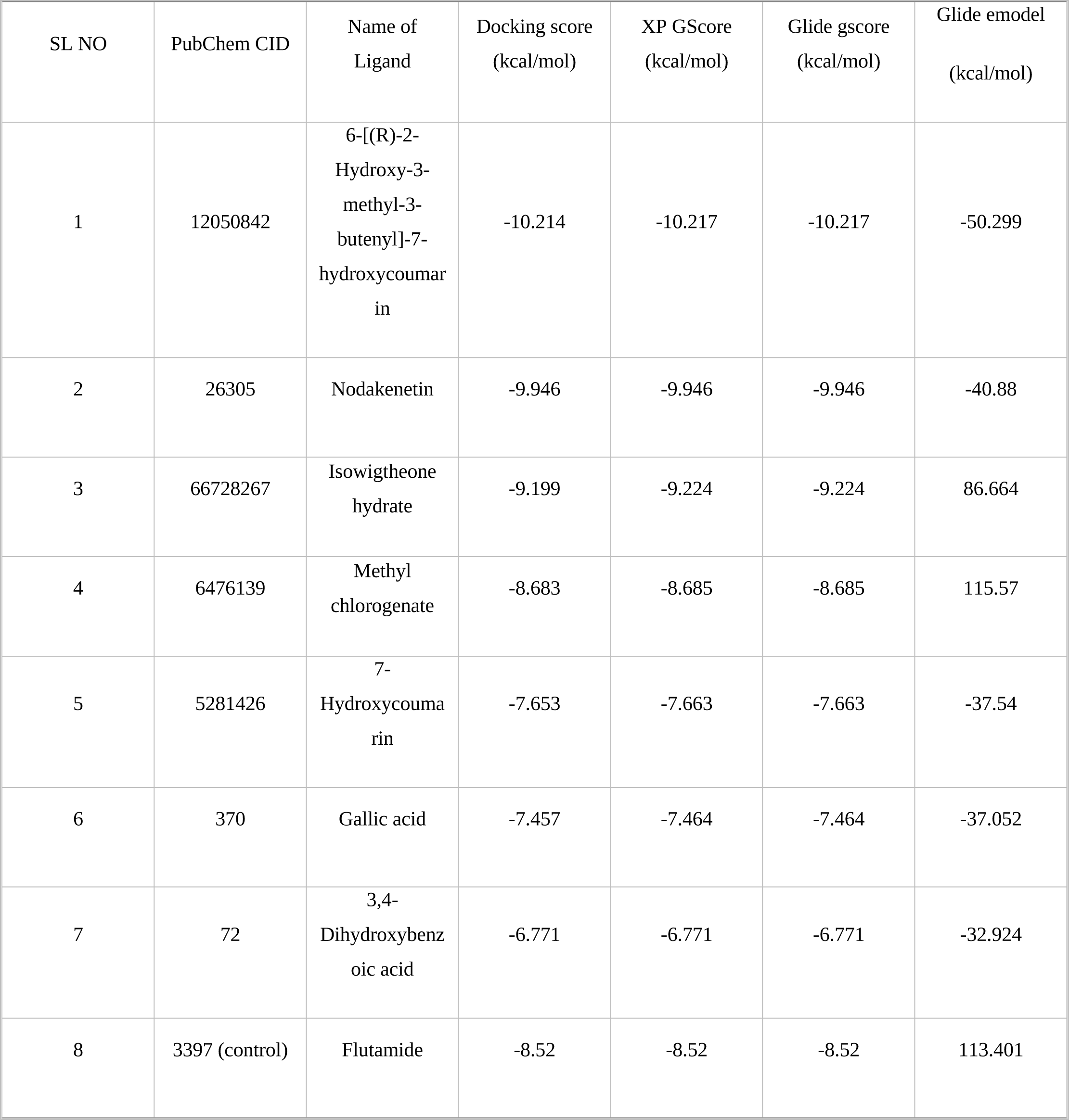
Site specific molecular docking score of all the phytocompounds with the targeted protein.

| SL NO | PubChem CID | Name of Ligand | Docking score (kcal/mol) | XP GScore (kcal/mol) | Glide gscore (kcal/mol) | Glide emodel (kcal/mol) |
| --- | --- | --- | --- | --- | --- | --- |
| 1 | 12050842 | 6-[(R)-2-Hydroxy-3-methyl-3-butenyl]-7-hydroxycoumarin | -10.214 | -10.217 | -10.217 | -50.299 |
| 2 | 26305 | Nodakenetin | -9.946 | -9.946 | -9.946 | -40.88 |
| 3 | 66728267 | Isowigtheone hydrate | -9.199 | -9.224 | -9.224 | 86.664 |
| 4 | 6476139 | Methyl chlorogenate | -8.683 | -8.685 | -8.685 | 115.57 |
| 5 | 5281426 | 7-Hydroxycoumarin | -7.653 | -7.663 | -7.663 | -37.54 |
| 6 | 370 | Gallic acid | -7.457 | -7.464 | -7.464 | -37.052 |
| 7 | 72 | 3,4-Dihydroxybenzoic acid | -6.771 | -6.771 | -6.771 | -32.924 |
| 8 | 3397 (control) | Flutamide | -8.52 | -8.52 | -8.52 | 113.401 |

### 3.3 Molecular Docking and Virtual Screening Analysis

Following the screening processes (PASS and ADMET property analysis), Glide docking with Extra precision was performed on all the 10 ligand phytochemicals (XP). Every plausible conformation for each low energy conformer in the specified binding site is determined using glide XP model ^34^. While the protein conformation is fixed, the torsional degrees of each ligand are relaxed during the process. Conversely, 7 out of the 10 phytochemicals show their docking score. The docking scores, Glide energy, emodel value, and the quantity of hydrogen bond forms were utilized to rank the preferred conformation for each ligand during the docking process. Considering the flexibility of both the ligand and the protein, it might be because of the Glide and Prime module that proper binding is depicted. Pose selection was facilitated by the emodel values. Analysis of the Glide energy term, which is Coulomb’s and van der Waal’s interaction, can also consider the energy of the ligand interaction. However, 6-[(R)-2-Hydroxy-3-methyl-3-butenyl]-7-hydroxycoumarin (CID 12050842) returned the best binding energies of -10.214 kcal/mol (table-3) but unfortunately, it has not directly involved with prostate cancer associated activities (results from pass online activities, table-1) that’s why we omitted this compound from the final selection list. Nodakenetin (CID 26305) gives -9.946 kcal/mol binding energy with the receptor protein and it is highly associated with prostate cancer as well as an androgen antagonist. However, another two compounds Isowigtheone hydrate (CID 66728267) and 7-Hydroxycoumarin (CID 5281426) gave binding energies of -9.199 kcal/mol and -7.653 kcal/mol, respectively. All these compounds are highly associated with prostate cancer treatment. But the compound 3,4-Dihydroxybenzoic acid (CID 72) has cancer associated activities (results from pass online activities, table-1) but its docking score is lower than others in the final docking result list (table-3), that’s why we did not consider this one as a high performing compound for final selected compound list. And from this screening process, the CID 26305, CID 66728267, and CID 5281426 were considered the best three phytochemicals for our further final analysis based on their docking score and for androgen antagonists. An FDA-approved antiandrogens drug flutamide (CID 3397) was considered as a reference compound for comparison with the three final selected phytochemicals.

### 3.4 Binding Interactions through Glide XP Docking

The selected three ligand interactions with the desired protein were analyzed using BIOVIA Discovery Studio Visualizer 21 tool. The Nodakenetin has been found to generate several conventional and carbon-hydrogen bonds with the targeted AR protein. It was found that two typical hydrogen bonds formed at the positions of Asn705 (3.22 Å) and Arg752 (6.31Å) (Figure 2). A Pi-Sigma and a Pi-Pi bond were also seen during the interaction in the position of Met745 (4.54 Å) and Phe764 (4.72Å), as well as five Pi-Alkyl bonds also found at the positions of Met749 (5.73 Å), Met780 (5.65 Å), Leu701 (5.06 Å), Leu704 (5.22 Å), and Leu880 (5.82 Å).

**Figure 2:**
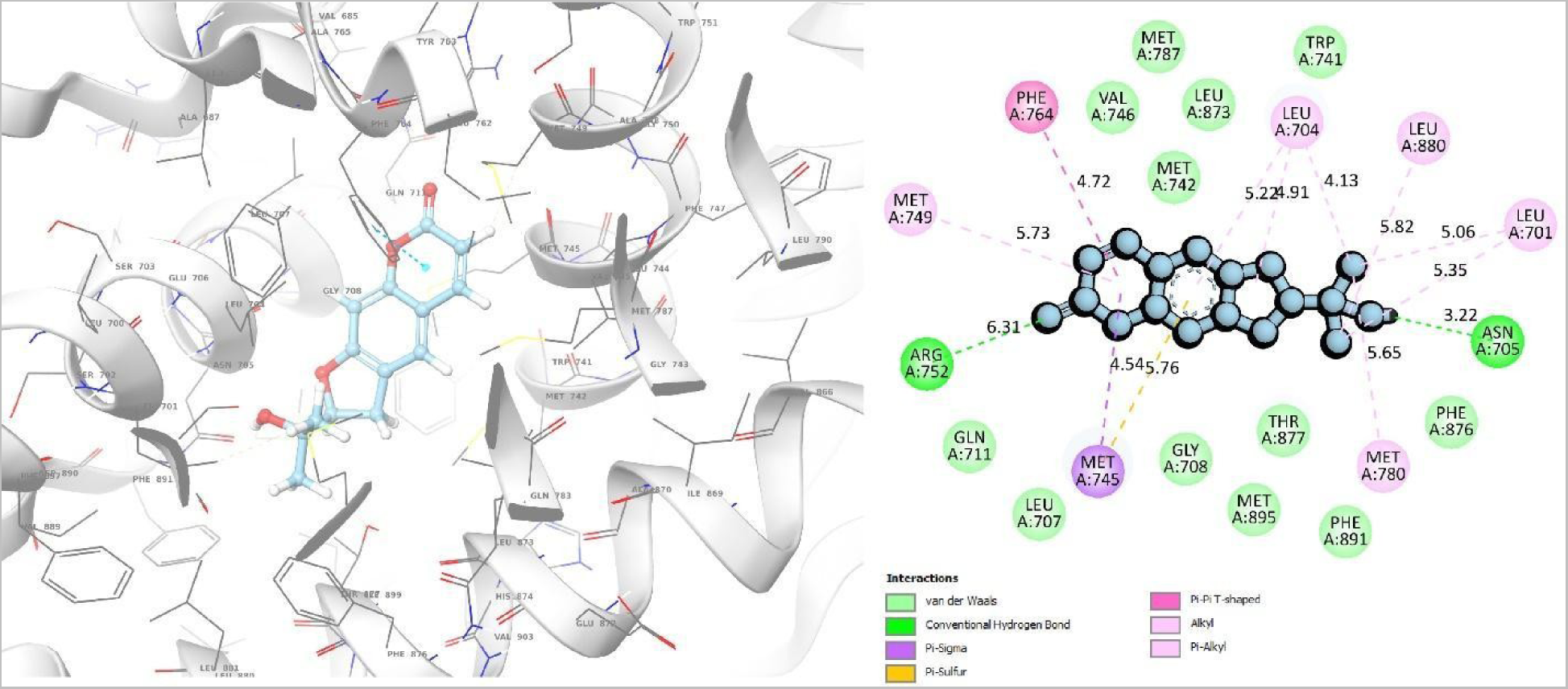
5T8E and Nodakenetin binding interactions. Left is the 3D interaction and right is the 2D interaction.

Three conventional hydrogen bonds were discovered in the Isowigtheone hydrate at the positions of Asn705 (3.81 Å), Arg752 (5.83 Å), and Met745 (3.83 Å) (Figure 3). There are five Pi-Alkyl bonds at Met780 (4.42 Å), Phe876 (3.85 Å), and Leu873 (4.04 Å). Additionally, a Pi-Pi link was discovered at Phe764 (5.26 Å). One conventional hydrogen bond, one Pi-Sulfur, one Pi-Alkyl, and one Pi-Pi bond were discovered in the interaction analysis of phytochemical 7-hydroxycoumarin at the positions of Leu704 (4.27 Å), Met787 (8.30 Å), Met745 (4.10, 5.41 Å), and Phe764 (5.34 Å) (Figure 4).

**Figure 3:**
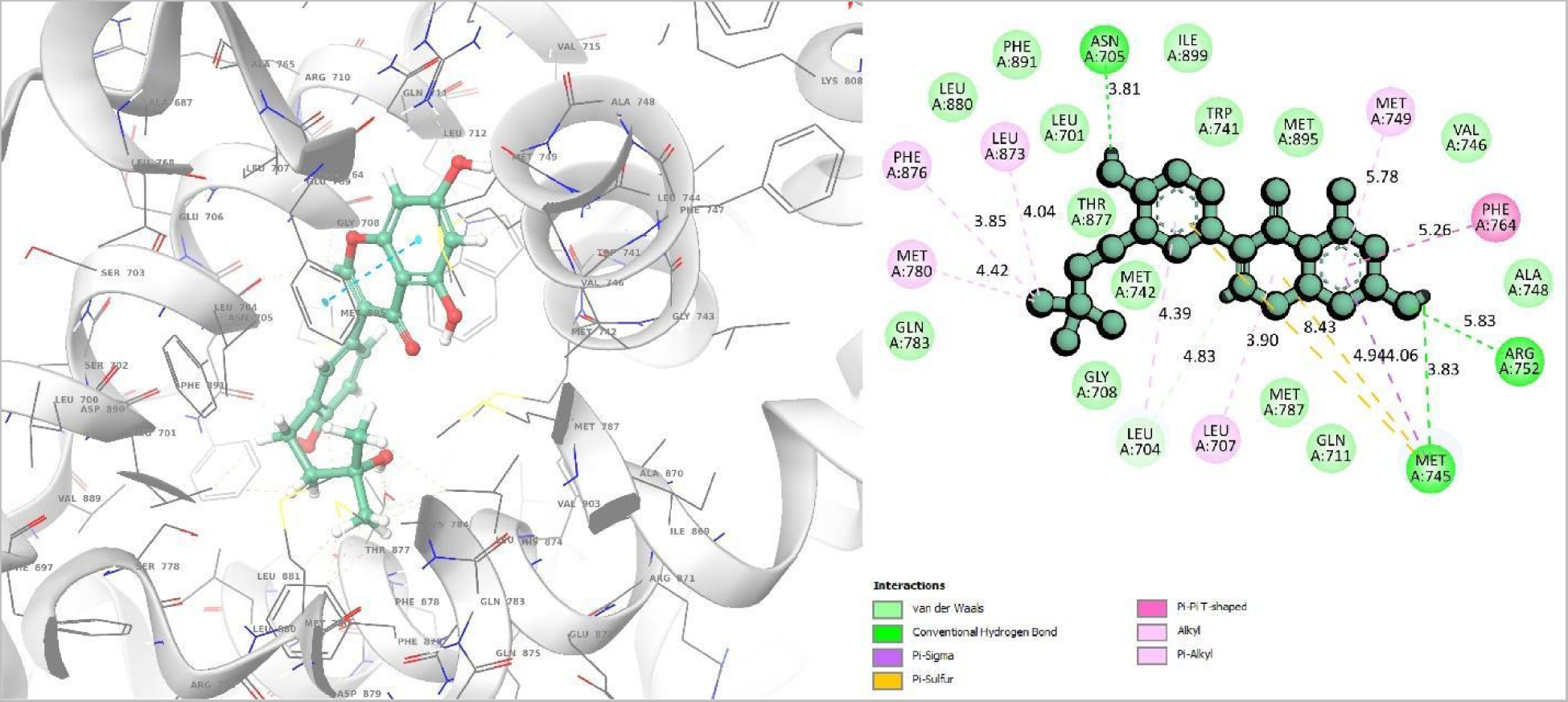
5T8E and Isowigtheone hydrate binding interactions. Left is the 3D interaction and right is the 2D interaction.

**Figure 4:**
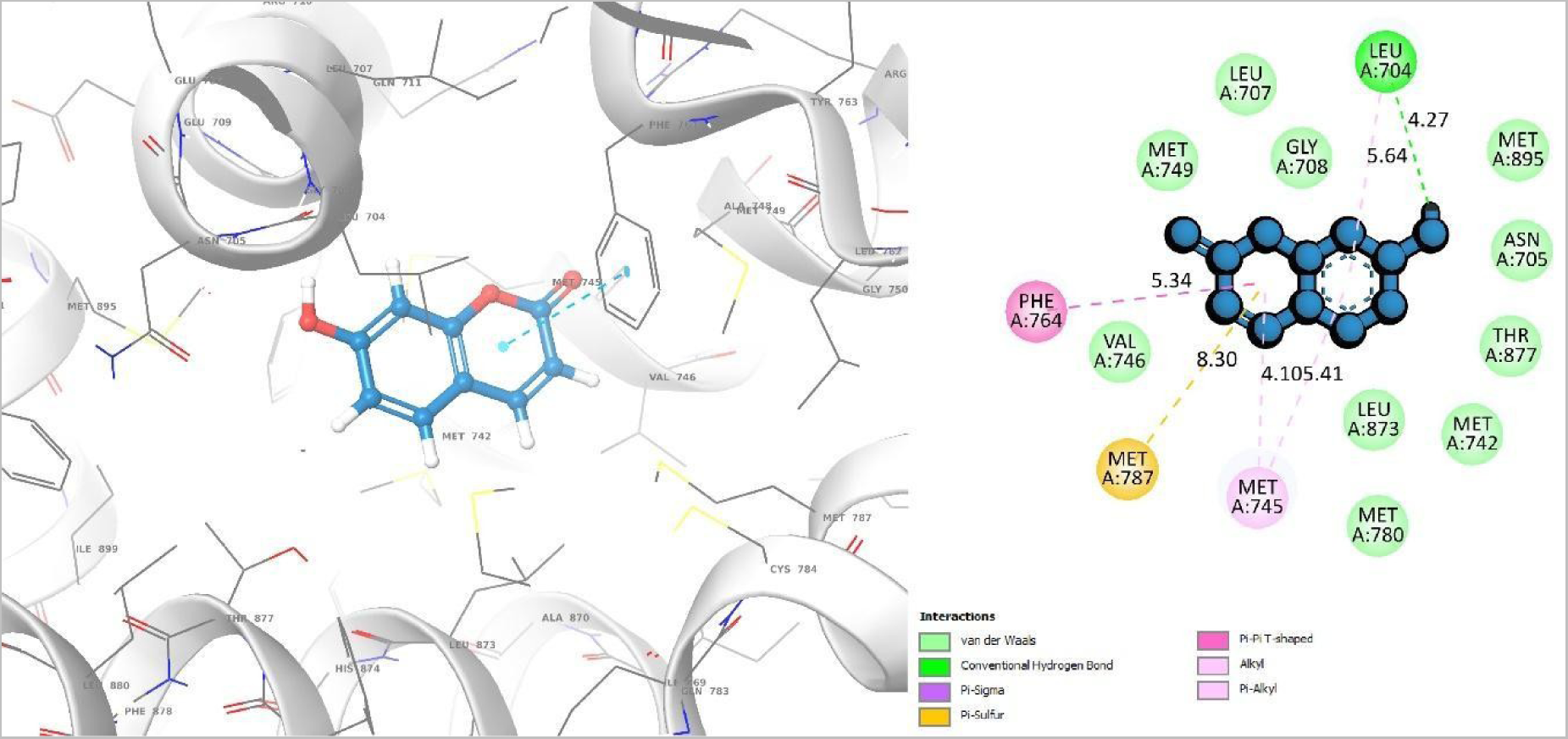
5T8E and 7-hydroxycoumarin binding interactions. Left is the 3D interaction and right is the 2D interaction.

### 3.5 MD Simulation Analysis

MD simulations are used to verify the stability of protein-ligand complexes in an artificial environment. Using MD simulations, one can track the changes in protein conformation over time to better understand how proteins move ^35^. Therefore, we performed 100 ns MD simulations of the protein-ligand complex structures to determine the candidate ligand molecules consistency in binding to the target receptor protein active site. The protein-ligands complex’s stability was compared using a food and drug administration (FDA) approved standard drug namely Flutamide (CID: 3397) which also shows a -8.52 kcal/mol docking score with the targeted receptor 5T8E. To analyze the MD simulation results, RMSD, RMSF, Rg, SASA, P-L contact, L-P contact, and hydrogen bond interactions were described.

#### RMSD Analysis of Protein

The average movement of atoms between two reference frames has been measured by using RMSD. The admissible range of the RMSD according to a reference frame is 1 to 3 Å ^36^. The RMSD progression of a protein (left Y-axis) and a ligand (right Y-axis) are shown in Figure 5. After all protein frames are aligned on the reference frame backbone, the RMSD for CID: 26305 (Nodakenetin), CID: 66728267 (Isowigtheone hydrate), CID: 5281426 (7-Hydroxycoumarin) and the control drug CID: 3397 (flutamide) is calculated (Figure 5). With the reference 5T8E protein backbone, all the three phytochemicals and the control drug performed very identical stable binding with the protein within a range of 0.6 - 2 Å (Figure 6).

**Figure 5:**
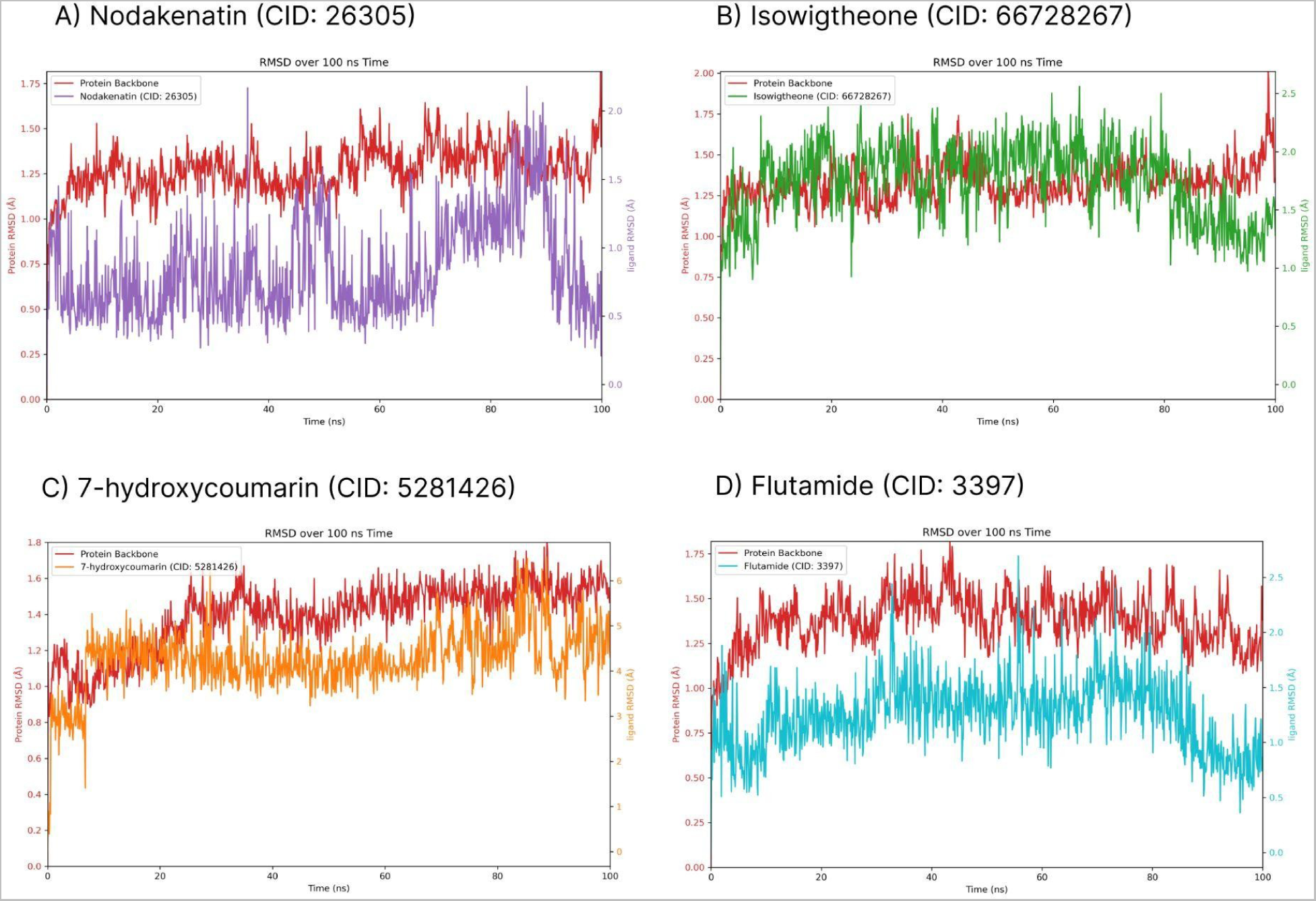
Protein-Ligand RMSD of Nodakenatin CID: 26305 (A), Isowigtheone CID: 66728267 (B), 7-hydroxycoumarin CID: 5281426 (C), Flutamide CID: 3397 (D), (E) Protein RMSD and f) Ligand RMSD of 100 ns simulation time.

**Figure 6:**
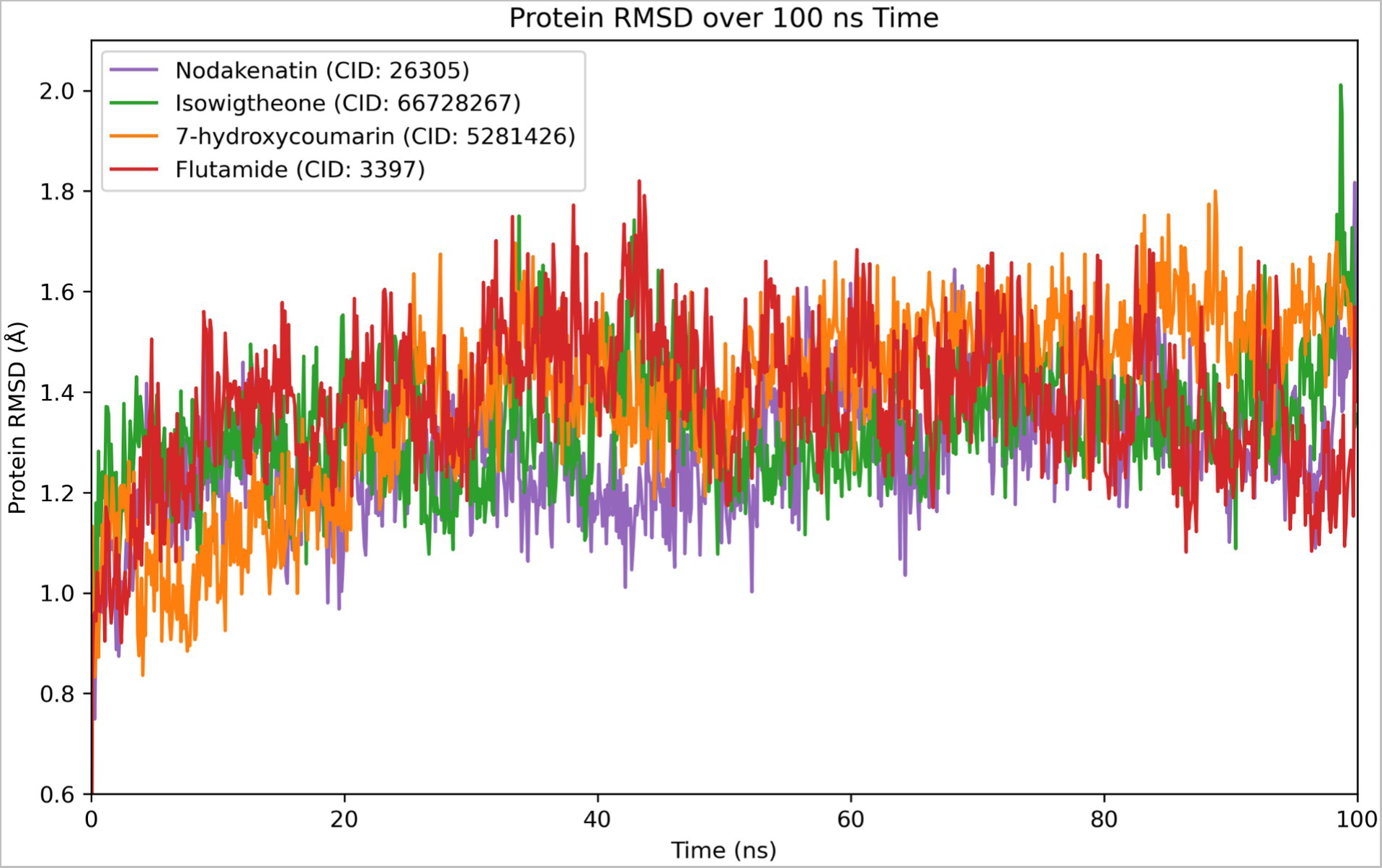
Protein RMSD over 100 ns simulation time. Nodakenatin CID: 26305-purple color, Isowigtheone CID: 66728267-green color, 7-hydroxycoumarin CID: 5281426-orange color, Flutamide CID: 3397 (Control drug)-red color.

The RMSD for the protein backbone is 0.6-2.0 Å for the 100 ns simulation runtime and it gives an average RMSD value of 0.8-1.4 Å from the first 20 ns (Figure 6). Also, the protein RMSD of Nodakenetin (CID: 26305) average value is 1.2-1.5 Å from 20-90 ns runtime. The same average value for the Isowigtheone hydrate (CID: 66728267). Also, the 7-Hydroxycoumarin (CID: 5281426) average RMSD value is 1.3-1.7 Å from 25-100 ns throughout the 100 ns runtime. The control drug average RMSD value is 1.2-1.8 Å through 20-80 ns of the whole simulation runtime.

It has been observed that the fluctuation was increasing for all phytochemicals at the first 40 ns. After that, the state of equilibration became apparent, and after 90 ns, the fluctuation started to decrease. Therefore, the RMSD helps to consider how the fluctuations will be optimized over the length of the extensive simulation runtime and how stable a ligand is regarding the protein and its binding pocket. The RMSD of the ligand is displayed in the plot after the protein-ligand complex is initially aligned on the reference protein backbone and the ligand RMSD heavy atoms are measured. Also, we computed the RMSD of the phytochemicals in that scenario to assess the stability of the phytochemicals compared to the control drug. In this instance, the binding complexes observations discovered optimal RMSD for the selected phytochemicals.

#### RMSD Analysis of Ligand

To assess the phytochemicals stability relative to the control drug, Ligand RMSD were calculated. Figure 7 shows the computed RMSD for the phytochemicals Nodakenetin (CID: 26305), Isowigtheone hydrate (CID: 66728267), 7-Hydroxycoumarin (CID: 5281426) and the control drug flutamide (CID: 3397). The complex docking structure was first aligned on the reference protein’s backbone to measure the RMSD of the phytochemicals. The phytochemicals can move away from their initial binding site because the observation found the optimum RMSD for two phytochemicals namely Nodakenetin (CID: 26305), Isowigtheone hydrate (CID: 66728267), except 7-Hydroxycoumarin (CID: 5281426), whose distance are significantly larger than the RMSD value of control drug flutamide (CID: 3397). It was observed that 7-Hydroxycoumarin (CID: 5281426) fluctuation was increasing at the start of 0 – 6 ns. After 6 ns, the state of equilibration became apparent, and after 66 ns, the fluctuation started to increase again, and the average deviation is 3.2-6 Å. Also, for Isowigtheone hydrate (CID: 66728267) average RMSD is 1.0-2.2 Å, for Nodakenetin (CID: 26305) the average RMSD is 1.0-2.4 Å that is compatible to the control drug. The control drug flutamide (CID: 3397) average RMSD value is 0.8-1.9 Å and they give stable fluctuations through the runtime of the 100 ns simulation runtime.

**Figure 7:**
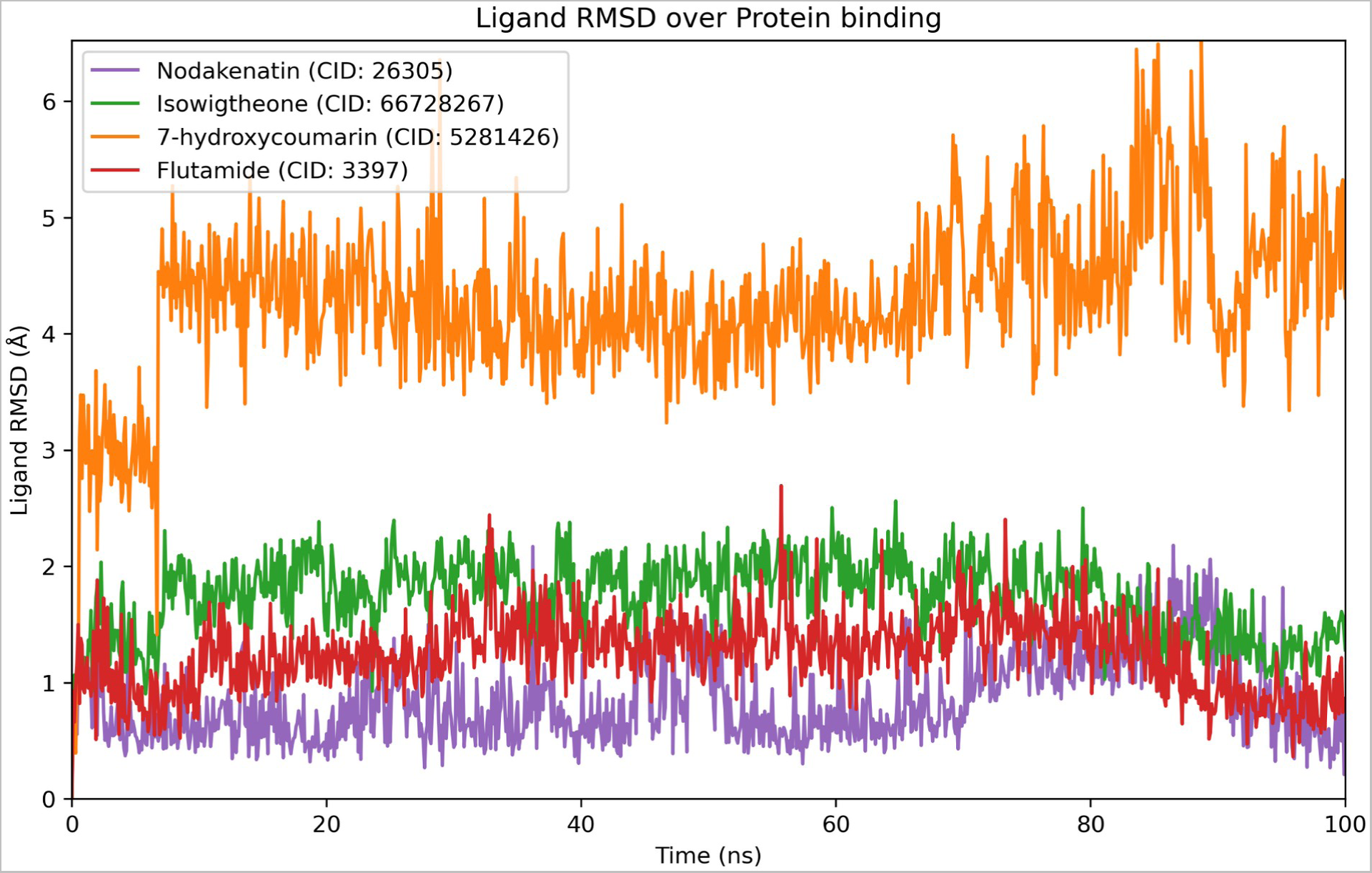
Ligand RMSD over protein binding during the 100 ns simulation time. Nodakenatin CID: 26305-purple color, Isowigtheone CID: 66728267-green color, 7-hydroxycoumarin CID: 5281426-orange color, Flutamide CID: 3397 (Control drug)-red color.

#### RMSF Analysis

The RMSF value is crucial for measuring the deviation of each atom’s position in a molecular dynamics simulation from its average position over the simulation period ^37^. The results are given in Figure 8, all the phytochemicals maintained an average fluctuation over the 100 ns simulation time. At the beginning of the simulation the 1-30 residue positions deviated to 2.1 Å. Then their average RMSF was 0.5-1.3 Å, except the position 170-178 atomic position. It is also noticeable that the 240-150 position increases the fluctuations. Nodakenetin (CID: 26305) maximum fluctuates in 20, 175 and 245 position but its average RMSF is 0.5 to 1.5 that is stable in comparison to the control drug. The control drug Flutamide (CID: 3397) demonstrated the greatest and most consistent fluctuations between 170 – 180 and 240-250 residual positions. The Isowigtheone hydrate (CID: 66728267) phytochemical fluctuates at a maximum of 4 Å in 245 and 2.6 Å in 175 residual positions. The RMSF of 7-Hydroxycoumarin (CID: 5281426), had the same flow as the control drug.

**Figure 8:**
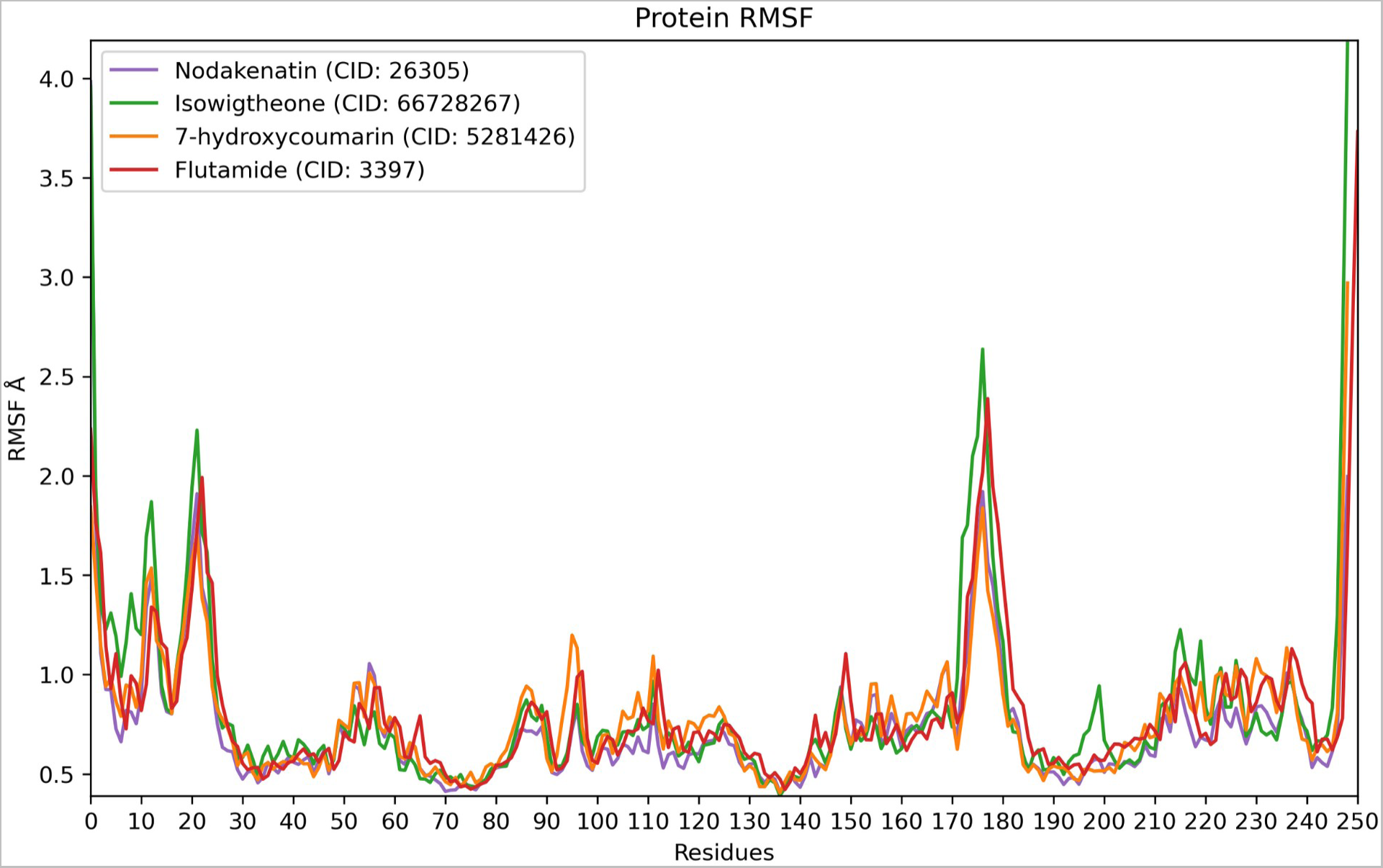
RMSF Analysis of 100 ns simulation time. Nodakenatin CID: 26305-purple color, Isowigtheone CID: 66728267-green color, 7-hydroxycoumarin CID: 5281426-orange color, Flutamide CID: 3397 (Control drug)-red color.

#### Protein-ligand Contacts Analysis

Throughout the 100 ns simulation, protein interactions with the ligand are observed. Figure 9 illustrates how these interactions can be divided into categories and summarized by type. The four types of protein-ligand interactions (or "contacts") appeared in the analysis including hydrogen bonds, hydrophobic interactions, ionic interactions, and water bridges. The subcategories of each interaction type were viewed using the desmond SID module. Throughout the trajectory time frame, the stacked bar charts are normalized. However, at the residue position of Asn705, which connects to other molecules via hydrogen and water bridge connections, the stacked bar charts for Nodakenetin (CID: 26305) shows an interaction fraction value (IFV) of 0.78 Å, which suggests that the particular interaction has been sustained for more than 70% of the simulated time (Figure 9A). The IFV found a maximum of 1.2 Å at Phe764 position that was produced by a hydrogen and water bridge bond as well as 0.98 Å at Asn705, which indicated that the Isowigtheone hydrate (CID: 66728267) phytochemical was maintaining the contacts over 100% and 90%, respectively (Figure 9B). Additionally, the phytochemical 7-Hydroxycoumarin (CID: 5281426) has an IFV value of 0.97 Å at the position of Asn705, which was created by a hydrogen and water bridge bond that was kept at a level of above 90% contact (Figure 9C). The control drug Flutamide (CID: 3397) was shown to form numerous contacts with the same residue within the same atom of the ligand (Figure 9D). However, the Nodakenetin (CID: 26305) shows 76% of interactions in the Asn705 residue from 0 to 100 ns of the simulation time. Also, the highest interactions for phytochemical 7-Hydroxycoumarin (CID: 5281426), are 42% with Leu704, 92% with Asn705 and 70% with residue Thr877 respectively.

**Figure 9:**
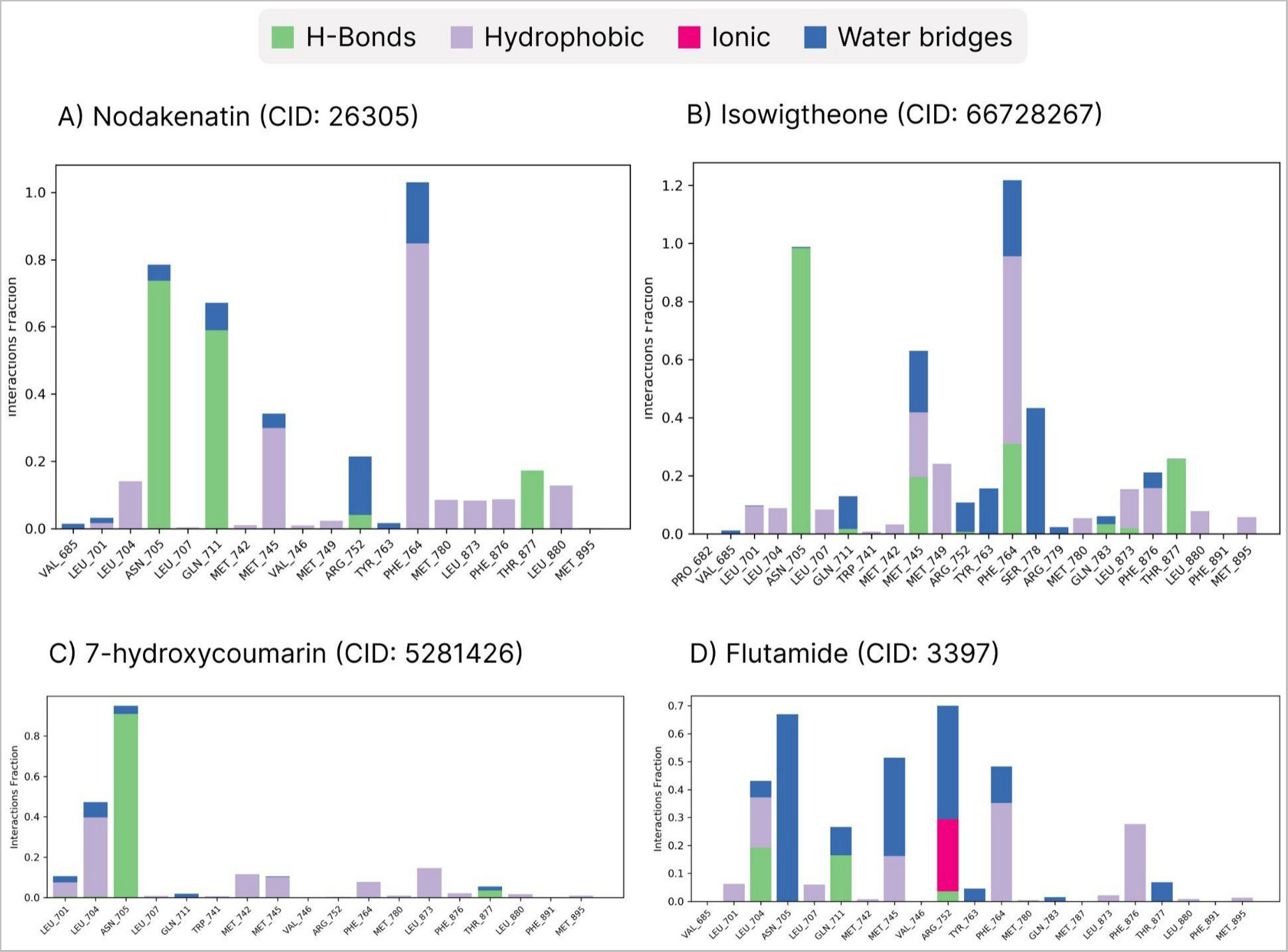
Protein-ligand contacts of (A) Nodakenatin CID: 26305, (B) Isowigtheone CID: 66728267, (C) 7-hydroxycoumarin CID: 5281426, (D) Flutamide CID: 3397 (Control drug).

#### Ligand Properties Analysis

**Figure 10:**
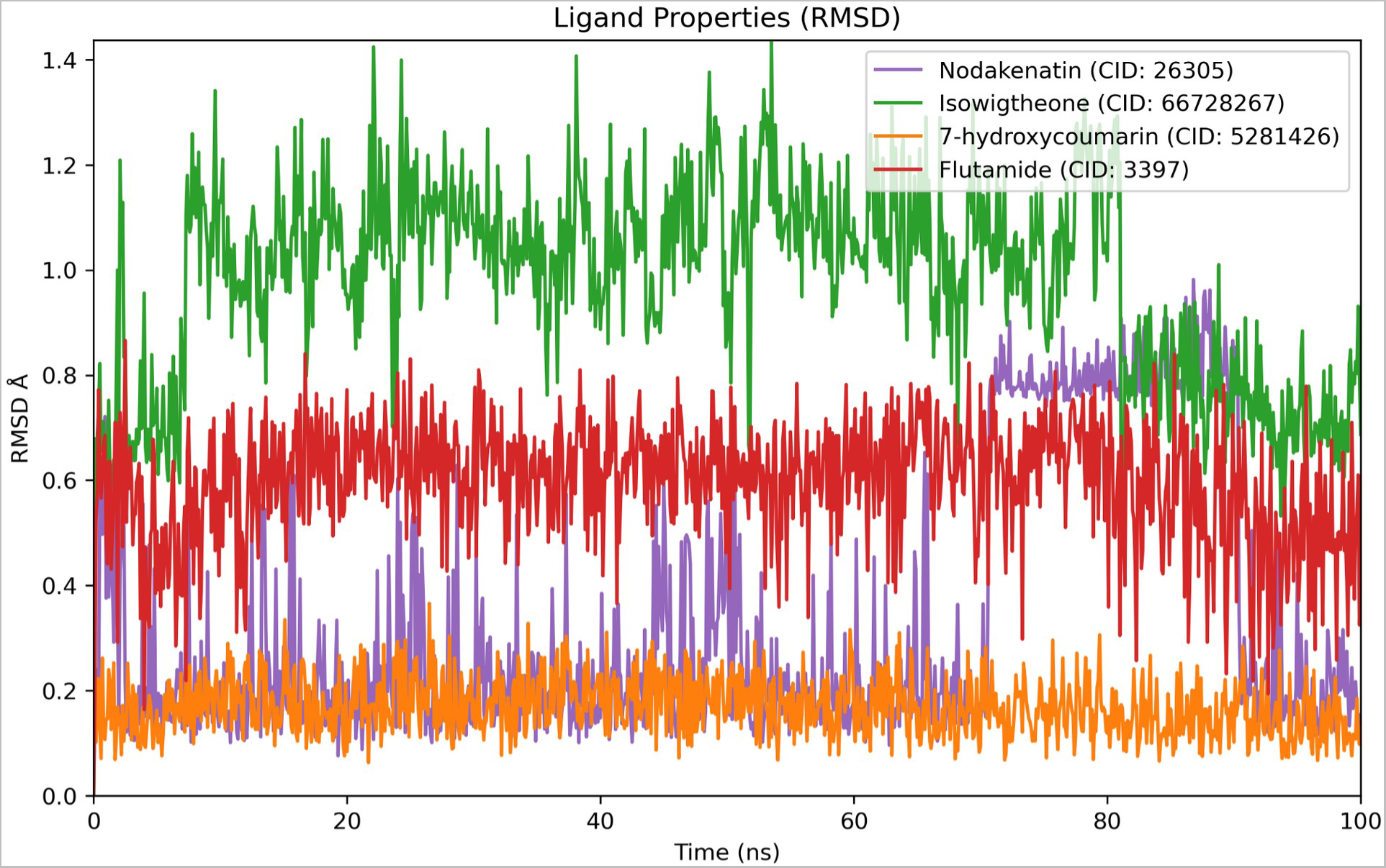
Ligand individual RMSD analysis during the simulation. (Nodakenatin CID: 26305-purple color, Isowigtheone CID: 66728267-green color, 7-hydroxycoumarin CID: 5281426-orange color, Flutamide CID: 3397 (Control drug)-red color.

#### Analysis of SASA, MolSA

Surfaces area accessible to solvents influences their construction as well as functionality underlying biologic biomolecules (SASA). The majority of about time, individual amino acid sequences of a protein’s surfaces function as activity domains may or may not combine with other components including receptors, which aids in understanding a molecule’s solvent-like characteristics (hydrophilic or hydrophobic) in addition to protein-ligand complexes. Biological macromolecules’ structural confirmation and functions are regulated by the amount of SASA. In most situations, amino acid residues on a protein’s surface function as active sites and/or interact with other molecules and ligands, which improves the understanding of a molecule’s solvent-like behavior (hydrophilic or hydrophobic) such as solubility, permeability, and bioavailability ^38, 39^. As a result of SASA value, the Isowigtheone hydrate (CID: 66728267) phytochemicals performed large surface areas during the 100 ns simulation time (Figure 11) indicating the high level of exposure of an amino acid residue to the selected compound in the complex systems. Nodakenetin (CID: 26305) and 7-Hydroxycoumarin (CID: 5281426) both show impressively stable receptor protein as well as control drug molecules (Figure 11).

**Figure 11:**
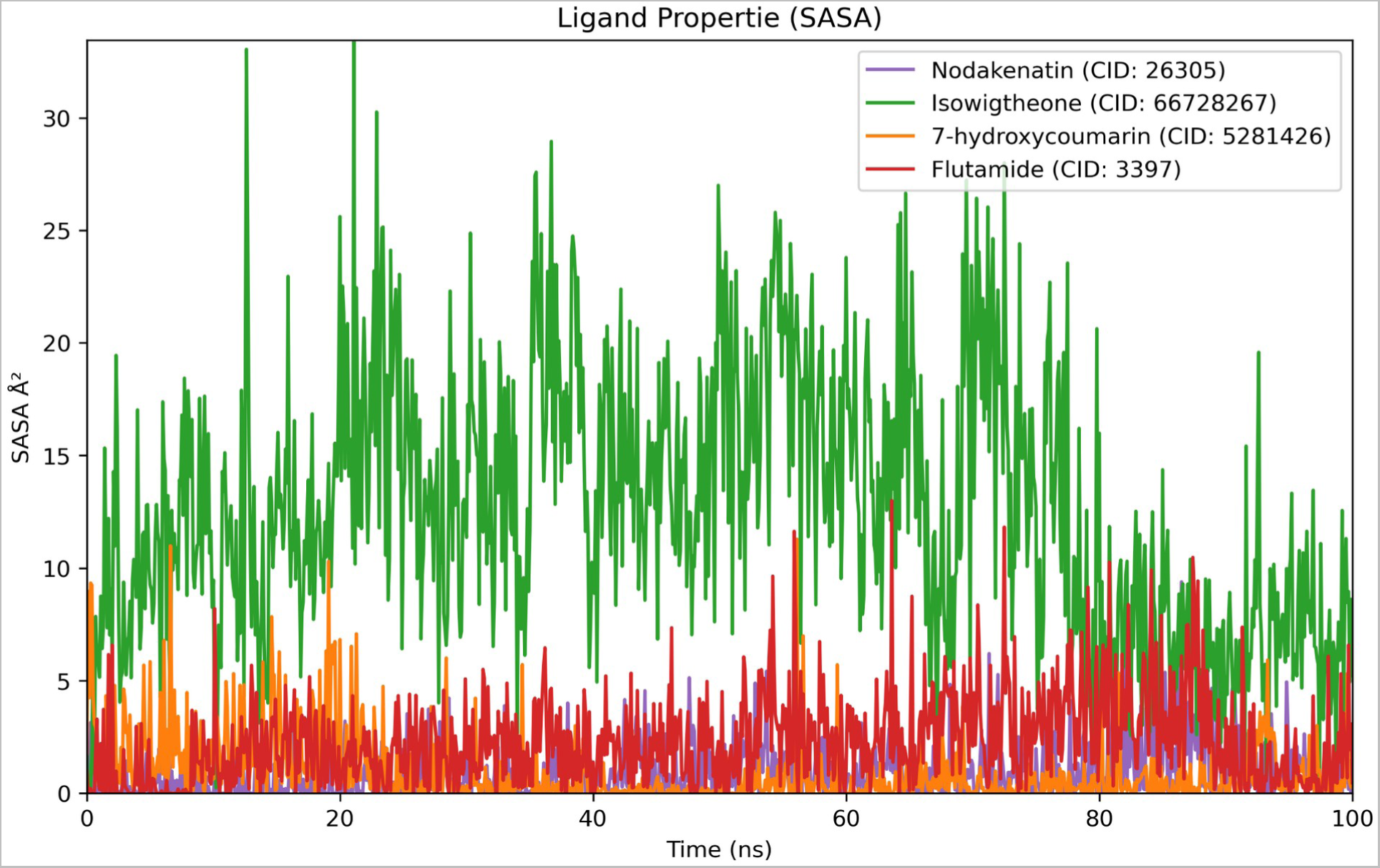
Ligand SASA analysis during the simulation. (Nodakenatin CID: 26305-purple color, Isowigtheone CID: 66728267-green color, 7-hydroxycoumarin CID: 5281426-orange color, Flutamide CID: 3397 (Control drug)-red color.

The MolSA is equivalent to a van der Waals surface area which is calculated with a 1.4 Å probe radius. In our *in-silico* study, Nodakenetin (CID: 26305), 7-Hydroxycoumarin (CID: 5281426) and Flutamide (CID: 3397) (Control drug) possessed the standard van der Waals surface area (Figure 12). Additionally, the compound Flutamide (CID: 3397) possessed a higher MolSA value compared to the other ligand compounds which effectively indicated the greater van der Waals surface area of the ligand compound.

**Figure 12:**
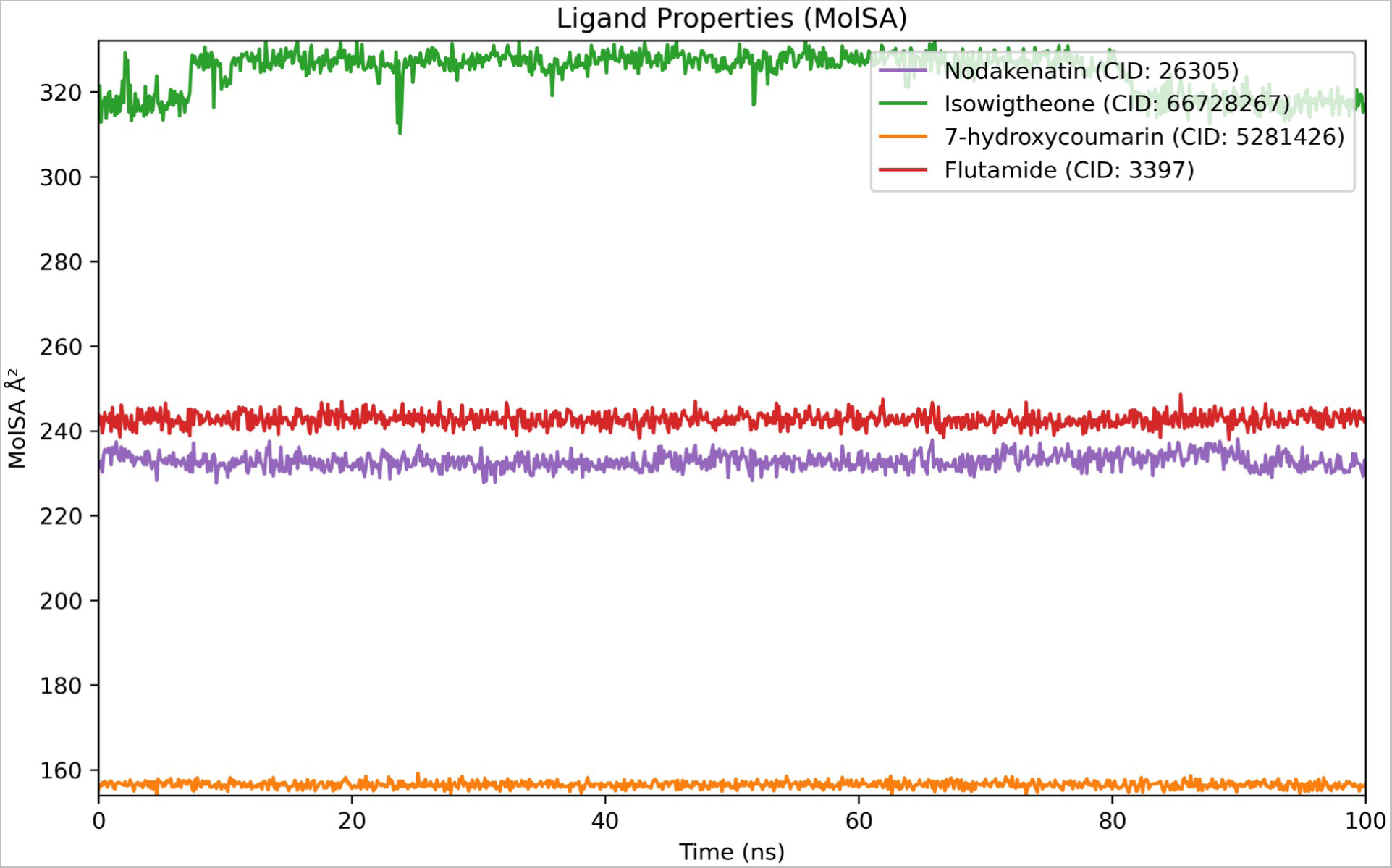
Ligand MolSA analysis during the simulation. (Nodakenatin CID: 26305-purple color, Isowigtheone CID: 66728267-green color, 7-hydroxycoumarin CID: 5281426-orange color, Flutamide CID: 3397 (Control drug)-red color.

#### H-bond Analysis

Hydrogen (H) bonds play a key role in both the formation and stabilization of protein structures and stabilize a ligand with the targeted protein in a ligand-protein complex ^40, 41^. In drug discovery study, H-bond influences the drug specificity, and it accelerates its metabolism and adsorption, therefore, the stability of the H-bond is crucial. In this study, we analyzed the receptor protein-phytochemical complexes H-bond stability during the 100 ns simulation runtime (Figure 13). Impressively, the results reveal that all the phytochemicals are maintaining H-bonding stability with the 5T8E androgen receptor at an optimized level during the 100 ns runtime. Isowigtheone hydrate (CID: 66728267) maintains larger hydrogen bonds at the beginning of the binding and its bonding in several atomic positions of 708-712, 744-746, 782, 771-779 protein residue numbers. Nodakenatin (CID: 26305) also has the H-bond in starting and the ending of residues. The control drug as well as the 7-hydroxycoumarin (CID: 5281426) does not maintain the H-bond interactions too long.

**Figure 13:**
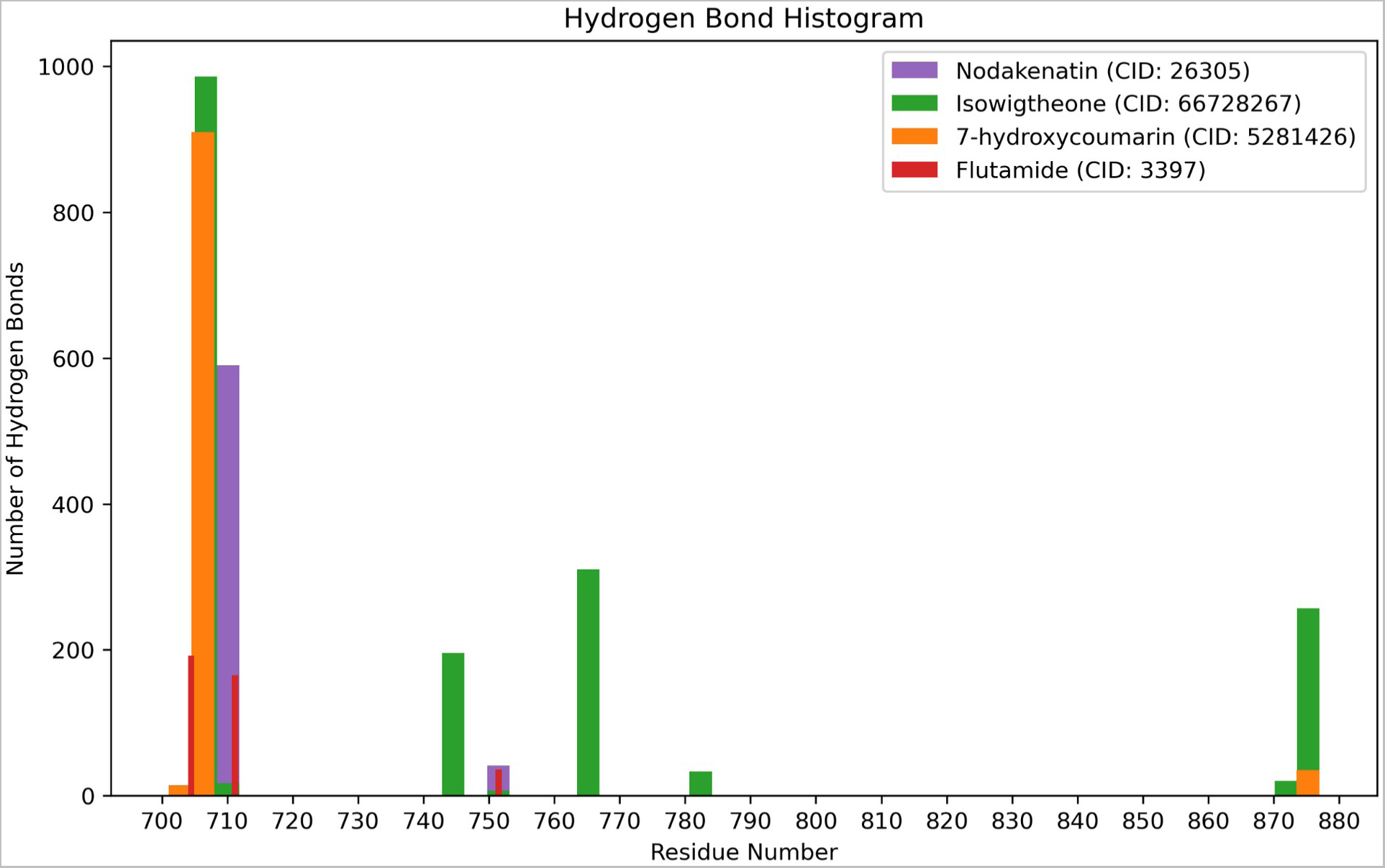
Hydrogen bond analysis result of Nodakenatin CID: 26305-purple color, Isowigtheone CID: 66728267-green color, 7-hydroxycoumarin CID: 5281426-orange color, Flutamide CID: 3397 (Control drug)-red color.

#### Post Simulation Binding Energy Analysis (MM/GBSA)

MMGBSA is known as the molecular mechanics-generalized Born surface area which can be performed to calculate ligand binding free energies and ligand strain energies for a set of ligands and a single receptor ^42^. Here, The greater the negative free energies of binding value (MMGBSA_dG_Bind_vdW for complex-receptor-ligand), the stronger the binding among the ligand compound with the targeted protein complex (Figure 14). Most importantly, the findings strongly depicted that the 7-hydroxycoumarin (CID: 5281426) with the receptor complex possessed strong binding free energy -49.65. In addition, the Isowigtheone (CID: 66728267) with the receptor complex exhibited binding free energy -44.64. However, the compounds Nodakenatin (CID: 26305) and Flutamide (CID: 3397) (Control drug) showed the lower binding free energy compared to the other 2 compounds at -33.63 and -32.97.

**Figure 14:**
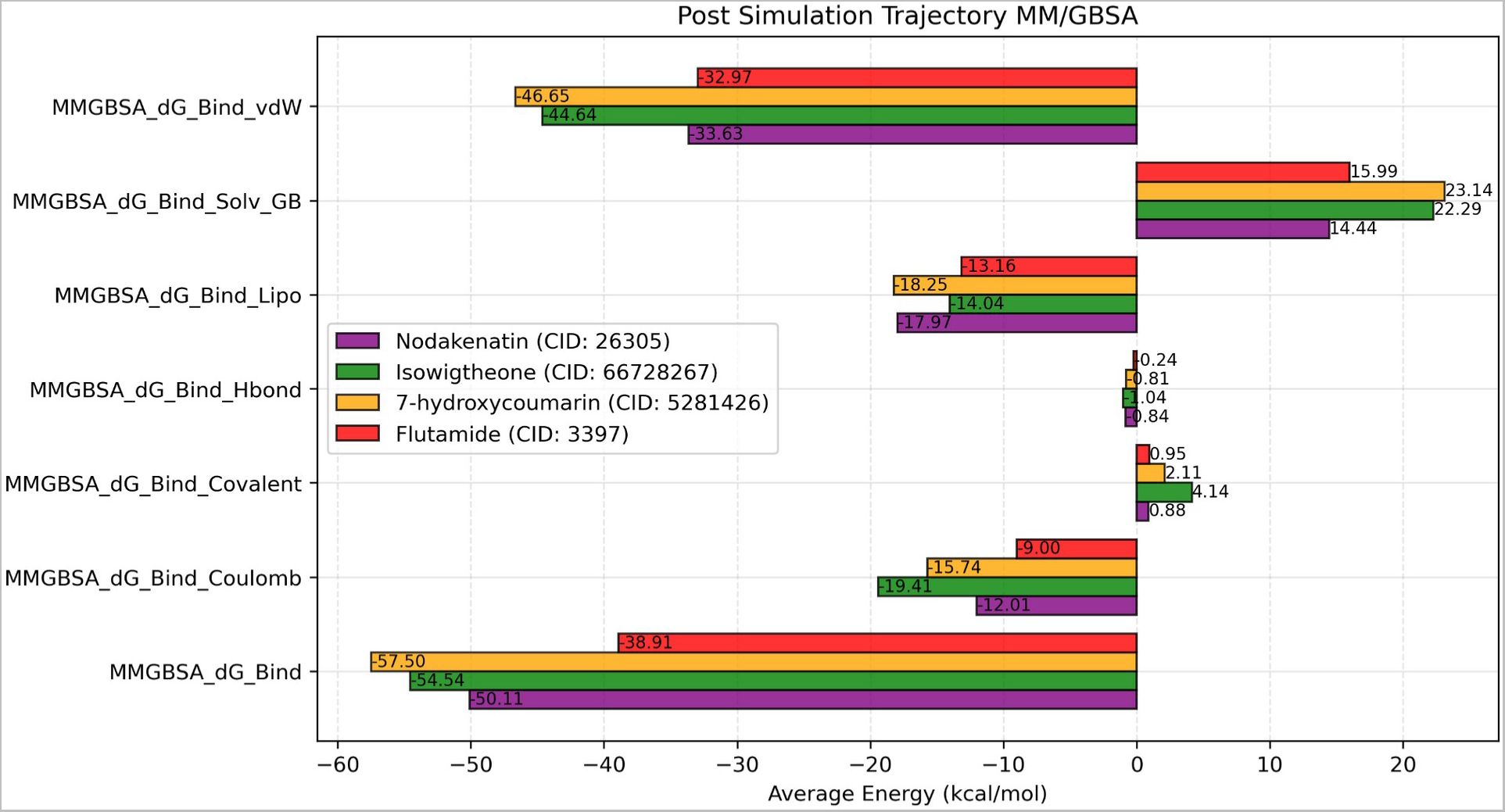
Post simulation trajectory MM/GBSA analysis of Nodakenatin CID: 26305-purple color, Isowigtheone CID: 66728267-green color, 7-hydroxycoumarin CID: 5281426-orange color, Flutamide CID: 3397 (Control drug)-red color.

## 4. Discussion

Prostate cancer is responsible for 1.4 million new cases of cancer in men and 375,000 deaths from the disease each year ^43, 44^. Androgen receptors (AR) play an important role in the development of prostate cancer ^45, 46^. Targeting the AR and determining the mechanisms of resistance to these agents continue to be central goals of drug development efforts ^46^. Whereas *Ficus hispida* has highly cytotoxic effects on prostate cancer cell lines as well as androgen receptors ^47^. In this study, we demonstrated that extracted phytochemicals from *Ficus hispida* fruits can interact with the AR of prostate cancer that can combat prostate cancer by targeting the AR 5T8E protein. The study aims to demonstrate that the phytochemicals in *Ficus hispida* can target AR and potentially serve as a novel drug for the treatment of prostate cancer. We screened a total of 89 phytochemicals of *Ficus hispida* that were extracted from different databases and literatures. This study focuses on the bioactive phytochemicals found in the fruits of *Ficus hispida* as it is popular as edible parts. Therefore only 13 bioactive phytochemicals were selected as potential ligands for the ADMET study. The ADMET study provides the most comprehensive data set for analyzing how the drug is processed by the human body ^48–50^. During this study, only 10 ligands passed the ADMET test based on the considered parameters of molecular weight < 500, donor hydrogen < 5, acceptor hydrogen <10, and violation of the rule of five < 3. Indeed, molecular docking was performed with the selected ligands against the receptor protein 5T8E.

In this study, prostate cancer inhibitors were screened computationally. Binding affinities of selected ligands with the targeted prostate androgen receptor are evaluated based on docking scores and intermolecular hydrogen bonding interactions. Thus, top-scoring phytochemicals had three hydrogen bonds with amino acids in the binding pocket.

Molecular docking aims to understand and predict molecular recognition, both structurally and energetically (i.e., predicting binding affinity). In biology, the hydrogen bond is essential for DNA structure and is usually considered by adding a term to the binding energy of a complex ^51, 52^. Based on the molecular docking results top seven ligands were selected considering their docking score (> -5) and the number of hydrogen bonds. The activity of the selected ligands was checked using PASS online activity prediction. Pa > Pi parameter was set for selecting the ligands activity and activities only related to the anti-prostate cancer were considered. Finally, the five ligands were selected for molecular dynamic simulation based on their docking score and hydrogen bond activities.

Molecular docking was carried out to evaluate the binding ability of the ligands to the 5T8E receptor protein. The docking score of CID: 26305 (Nodakenetin) was -9.946 kcal/mol and bound to the 5T8E receptor with two conventional hydrogen bonds in ASN 705 and ARG 752 residues. CID: 6476139 (Methyl chlorogenate) bound to the receptor with five conventional hydrogen bonds and docking score was -8.685 kcal/mol. CID: 66728267 (Isowigtheone hydrate) exhibited -9.199 kcal/mol and three conventional hydrogen bonds. Both CID: 370 (Gallic acid) and CID: 5281426 (7-Hydroxycoumarin) bound to the receptor with the single conventional hydrogen bond and scored -7.457 kcal/mol and -7.653 kcal/mol respectively in molecular docking. All these five ligands were found to have prostate cancer treatment activity in PASS online ligand activity prediction, though gallic acid and 7-hydroxycoumarin had lower Pa value (< 0.3). Another compound, 6-[(R)-2-Hydroxy-3-methyl-3-butenyl]-7-hydroxycoumarin scored better than other ligands in molecular docking but it was not considered for dynamic simulation for the lack of any anti-prostate cancer activity. The data of molecular docking indicates that the binding affinity of the selected ligands was higher, and the PASS online prediction confirms their possible potential against prostate cancer.

MD simulation explores protein-ligand binding stability ^53^. The MD simulation also provides intermolecular interaction information ^41, 54^. Based on RMSD, simulation equilibrium can be determined. On the contrary, fluctuations between 1–3 Å in a reference protein structure are acceptable, but larger values indicate large conformational change and instability ^16^. The protein RMSD data depicts that all the selected ligands performed well, and their average fluctuation forms the frame of the protein backbone is 0.5 Å. The fluctuation from 1-3 Å is acceptable, and the range of more than 3 Å is considered unstable. To check the stability of the ligands the ligand RMSD was calculated along with the control compound. In ligand RMSD CID: 26305 (Nodakenetin) and CID: 66728267 (Isowigtheone hydrate) showed more stability than other phytochemicals. Compared to the control compound, CID: 6476139 (Methyl chlorogenate), CID: 5281426 (7-Hydroxycoumarin), and CID: 370 (Gallic acid) exceeded the acceptable fluctuation range (3 Å). Also, the ligand properties analysis specially the H-bond analysis revealed that the *Ficus hispida* fruits phytochemicals are able for drug adsorption, metabolism, and specificity.

In an *in vitro* and *in vivo* study, Nodakenetin inhibited leukemia cell growth ^55^. Isowigtheone hydrate may also be effective as an anticancer agent ^56, 57^. Methyl chlorogenate could function as a therapeutic agent against benign prostatic hypertrophy ^58^. In different *in vitro* studies, 7-hydroxycoumarin exhibited anti-cancer potentials by inhibiting malignant cell growth and disturbing the cell cycle ^59–61^. However, studies on the anti-prostate cancer properties of these phytochemicals are unavailable. In contrast, gallic acid is an effective anti-prostate cancer agent. In several *in vitro* and *in vivo* studies, by inducing cell death, provoking DNA damage, and inhibiting histone deacetylase (HDACs) ^62–64^.

According to similar androgen receptor (PDB ID: 5T8E) and anti-prostate cancer therapeutic studies, researchers suggested that the androgen receptor is a promising target for prostate cancer treatment ^9, 65–67^. Although no *in vitro* or *in vivo* studies have been performed for the treatment of prostate cancer with *Ficus hispida*, so this in silico study may serve as a starting point for future research. For instance, the *Ficus hispida* fruits selected phytochemicals CID: 26305 (Nodakenetin), CID: 66728267 (Isowigtheone hydrate), CID: 6476139 (Methyl chlorogenate), CID: 5281426 (7-Hydroxycoumarin) and CID: 370 (Gallic acid) have lead drug specificity. However, we had a limitation for the phytochemical activity prediction that the PASS online prediction score is not > 0.5. To find out whether these phytochemicals have the potential to fight prostate cancer, additional *in vitro* or *in vivo* studies are needed. Therefore, further research into it may someday lead to a significant treatment for prostate cancer.

## 5. Conclusion

Androgen receptor plays a significant role in the progression of prostate cancer and therefore is a known target for the development of anticancer agents against prostate cancer. In this study, phytochemicals of *Ficus hispida* fruit were explored for potential drug-like compounds that can target AR. PASS online and ADMET tools screened a total of 10 phytochemicals based on their drug-like properties. binding affinity to AR. Of these, a total of three phytochemicals, including nodakenetin, isowigtheone hydrate, and 7-Hydroxycoumarin were selected by docking score. The binding stability of these three phytochemicals with AR was confirmed by MD simulation, suggesting that these compounds can be exploited for future anticancer drug development. However, further validation through in vitro and in vivo experimental approaches is required to develop *Ficus hispida* fruit phytochemicals as potential anticancer therapy against prostate cancer.

### Author Contributions

M.H.R. designed the project, generated the data, analyzed the data, and wrote the manuscript. M.S.Z. and P.B. analyzed the data and wrote the manuscript. S.I., R.A.I.R., and A.A.M.S. wrote the manuscript. B.G. and R.I. collected the data. M.J.U., M.A.H., M.A.R., W.K and B.K. supervised and reviewed the manuscript. All authors read and approved the final manuscript.

## Acknowledgments

We express our gratitude to the ABEx Bio-Research Center for providing the necessary software, hardware, and bioinformatics lab support (https://research.abexbio.com/).

