## Supplemental Table 1 for "Identification of Therapeutic Leads from *Ficus hispida* Fruit Phytochemicals against Prostate Cancer Using Pharmacoinformatic and Molecular Dynamics Simulation Approach"

**Table 2:** ADMET result

| SL NO | CID/SID | Phytochemicals | mol_MW | SASA | Volume | QPlogPw | QPlogPo/w | QPlogS | QPPCaco | QPlogKp | QPlogKhsa | RuleOfFive |
|---|---|---|---|---|---|---|---|---|---|---|---|---|
| 1 | 72 | 3,4-Dihydroxybenzoic acid | 154.122 | 329.839 | 502.685 | 9.936 | 0.014 | -0.779 | 27.436 | -4.61 | -0.905 | 0 |
| 2 | 370 | Gallic acid | 170.121 | 340.445 | 523.341 | 12.016 | -0.585 | -0.681 | 10.027 | -5.488 | -0.987 | 0 |
| 3 | 26305 | Nodakenetin / (-)-Marmesin | 246.262 | 466.594 | 789.661 | 7.69 | 2.048 | -3.094 | 984.592 | -2.672 | -0.057 | 0 |
| 4 | 5281426 | 7-Hydroxycoumarin | 456.707 | 683.818 | 1387.897 | 8.06 | 6.2 | -6.712 | 342.203 | -2.829 | 1.331 | 1 |
| 5 | 5490139 | Alpinumisoflavone | 162.145 | 344.677 | 536.547 | 7.338 | 0.71 | -1.425 | 624.506 | -2.988 | -0.512 | 0 |
| 6 | 6476139 | Methyl chlorogenate / | 336.343 | 589.7 | 1026.72 | 8.613 | 3.673 | -5.174 | 687.613 | -2.371 | 0.559 | 0 |

| | | | | | | | | | | | | |
|---|---|---|---|---|---|---|---|---|---|---|---|---|
| | | Chlorogenic acid methyl ester | | | | | | | | | | |
| 7 | 12050842 | 6-[(R)-2-Hydroxy-3-methyl-3-butenyl]-7-hydroxycoumarin | 368.34 | 643.055 | 1111.512 | 18.784 | 0.045 | -3.069 | 20.553 | -5.271 | -0.63 | 0 |
| 8 | 44570408 | (6R,9R)-Roseoside | 246.262 | 482.857 | 817.506 | 9.861 | 1.484 | -2.608 | 421.722 | -3.033 | -0.286 | 0 |
| 9 | 66728267 | Isowigtheone hydrate | 356.374 | 622.184 | 1098.6 | 11.292 | 2.826 | -4.396 | 125.1 | -3.586 | 0.253 | 0 |
| 10 | 101416188 | Murrayaculatine | 207.185 | 358.098 | 606.273 | 6.486 | -0.919 | -0.825 | 20.622 | -5.613 | -0.534 | 0 |
