## Supplemental Table 2 for "Identification of Therapeutic Leads from *Ficus hispida* Fruit Phytochemicals against Prostate Cancer Using Pharmacoinformatic and Molecular Dynamics Simulation Approach"

**Table 1:** PASS online prediction results

| SL No. | CID/SID | Phytochemicals | Pa | Pi | Activity |
|---|---|---|---|---|---|
| 1. | 72 | 3,4-Dihydroxybenzoic acid | 0.262 | 0.100 | Prostate disorders treatment |
| | | | 0.383 | 0.020 | Cancer-associated disorders treatment |
| | | | 0.170 | 0.044 | Cancer procoagulant inhibitor |
| 2. | 370 | Gallic acid | 0.183 | 0.085 | Prostate cancer treatment |
| | | | 0.198 | 0.160 | Prostate disorders treatment |
| | | | 0.374 | 0.024 | Cancer-associated disorders treatment |
| | | | 0.167 | 0.045 | Cancer procoagulant inhibitor |
| 3. | 26305 | Nodakenetin / (-)-Marmesin | 0.072 | 0.034 | Androgen antagonist |
| | | | 0.302 | 0.041 | Prostate cancer treatment |
| | | | 0.284 | 0.085 | Prostate disorders treatment |
| | | | 0.085 | 0.028 | Androgen antagonist |
| | | | 0.475 | 0.013 | Prostate cancer treatment |
| | | | 0.456 | 0.027 | Prostate disorders treatment |
| 5. | 5281426 | 7-Hydroxycoumarin | 0.099 | 0.023 | Androgen antagonist |
| | | | 0.359 | 0.050 | Prostate disorders treatment |

| | | | | | |
|---|---|---|---|---|---|
| | | | 0.283 | 0.046 | Prostate cancer treatment |
| | | | 0.334 | 0.055 | Cancer-associated disorders treatment |
| 6. | 5490139 | Alpinumisoflavone | 0.253 | 0.055 | Prostate cancer treatment |
| | | | 0.227 | 0.208 | Cancer-associated disorders treatment |
| | | | 0.075 | 0.032 | Androgen antagonist |
| | | | 0.615 | 0.012 | Prostate disorders treatment |
| | | | 0.212 | 0.069 | Prostate cancer treatment |
| 8. | 6476139 | Methyl chlorogenate / Chlorogenic acid methyl ester | 0.317 | 0.037 | Prostate cancer treatment |
| 9. | 12050842 | 6-[(R)-2-Hydroxy-3-methyl-3-butenyl]-7-hydroxycoumarin | 0.227 | 0.207 | Cancer-associated disorders treatment |
| 10. | 44570408 | (6R,9R)-Roseoside | 0.239 | 0.121 | Antineoplastic (pancreatic cancer) |
| 11. | 66728267 | Isowigtheone hydrate | 0.095 | 0.024 | Androgen antagonist |
| | | | 0.429 | 0.018 | Prostate cancer treatment |
| 12. | 101416188 | Murrayaculatine | 0.370 | 0.026 | Cancer-associated disorders treatment |
| | | | 0.118 | 0.089 | Cancer procoagulant inhibitor |
| 13. | 463911958 | Benzyl alpha-beta-glucopyranoside | 0.340 | 0.035 | Antineoplastic (pancreatic cancer) |
